# *In vivo* identification and validation of novel potential predictors for human cardiovascular diseases

**DOI:** 10.1101/2021.02.03.429563

**Authors:** Omar T. Hammouda, Meng Yue Wu, Verena Kaul, Jakob Gierten, Thomas Thumberger, Joachim Wittbrodt

## Abstract

Genetics crucially contributes to cardiovascular diseases (CVDs), the global leading cause of death. Since the majority of CVDs can be prevented by early intervention there is a high demand for predictive markers. While genome wide association studies (GWAS) correlate genes and CVDs after diagnosis and provide a valuable resource for such markers, preferentially those with previously known or suspected function are addressed further. To tackle the unaddressed blind spot of understudied genes, we particularly focused on the validation of heart GWAS candidates with little or no apparent connection to cardiac function. Building on the high conservation of basic heart function and underlying genetics from fish to human we combined CRISPR/Cas9 genome editing of the orthologs of human GWAS candidates in isogenic medaka with automated high-throughput heart rate analysis. Our functional analyses of understudied human candidates uncovered a prominent fraction of heart rate associated genes from adult human patients displaying a heart rate effect in embryonic medaka already in the injected generation. Following this pipeline, we identified 16 GWAS candidates with potential diagnostic and predictive power for human CVDs.

## Introduction

Genetics crucially contributes to the development and progression of cardiovascular diseases (CVDs), the global leading cause of death (Kathiresan & Srivastava 2012; Cambien & Tiret 2007). Elevated resting heart rate in humans has been widely considered as a potential, modifiable risk factor of cardiovascular and all-cause mortality (Beere et al. 1984; DYER et al. 1980; MD et al. 2010; Gillum et al. 1991). Since the majority of CVDs can be prevented by early intervention (McGill et al. 2008) there is a high demand for diagnostic and predictive CVD markers. Genome wide association studies (GWAS) on human patients correlate genes and CVDs after diagnosis and provide a valuable resource for those putative markers (Eicher et al. 2015). However, genes with pre-assigned cardiac functions are usually more likely to be addressed further, while uncharacterized genes or those with no pre-existing evidence to the heart are often neglected. This is likely due to the lack of experimental pipelines for the rapid and robust validation of such markers with implications for heart function. Recently, we demonstrated the power of targeted genome editing in the small animal model system medaka *(Oryzias latipes)* to validate trabeculation-associated genes (Meyer et al. 2020). The ease of manipulation combined with robust acquisition and analysis pipelines highlight the power of using fish embryos in high-throughput applications (Gierten et al. 2020; Shankaran et al. 2018; Lessman 2011; Oxendine et al. 2006). Embryos of fish model systems undergo extrauterine development in a transparent egg. This allows to monitor heart development and heart rate non-invasively in live undisturbed embryos for an extended period of time. Heart development, function and physiology in fish, though simpler, is comparable to mammals (Nemtsas et al. 2010; Yonekura et al. 2018; Gut et al. 2017). Here we combined targeted genome editing via CRISPR/Cas9 (Stemmer et al. 2015; Thumberger et al. 2021) with automated high-throughput imaging and heart rate analysis in isogenic medaka embryos (Gierten et al. 2020) to enable functional analyses directly in the injected generation (F0). We tested the performance our assay with a positive control *(nkx2-5),* evaluated the random discovery rate and analyzed 40 heart associated genes identified from human GWAS. Our assay uncovered that 57% of candidates assigned to human heart rate in GWAS also affected heart rate in fish embryos. We have thus experimentally validated understudied human GWAS candidates, identifying 16 genes with potential diagnostic and predictive power for human CVDs.

## Results

For the straight forward functional validation of GWAS candidates we aimed at combining CRISPR/Cas9-mediated targeted gene inactivation in the injected generation of medaka embryos with high content screening approaches to validate the impact of the loss-of-function on the heart rate (Fig 1A). We first assessed the efficiency of the CRISPR/Cas9 system by targeting the *green fluorescent protein (gfp)* gene in a transgenic medaka reporter line expressing GFP and mCherry fluorescent proteins exclusively in the heart via the *cardiac myosin light chain 2 (cmlc2)* promoter. Injection of the *heiCas9* mRNA, i.e. a Cas9 equipped with an early active nuclear localization signal (Thumberger et al. 2021), together with a guide RNA targeting *gfp* into medaka embryos at the 1-cell stage resulted in a complete loss of GFP expression in the heart (n=8/8) (Fig 1B and Fig S1). Only when injected into a single blastomere of the four-cell stage or later, a mosaic pattern was observable. This demonstrates the high efficiency of our approach already in the injected generation.

For our functional validation assay, we used the cardiac-specific homeobox-containing transcription factor NKX2-5 as a positive control. In human patients, a single amino acid mutation in the homeodomain (R141C) was previously associated with atrial septal defect (ASD) and shown to cause delayed heart morphogenesis in adult mice (Zakariyah et al. 2017). To test our high-throughput imaging and heart rate analysis pipeline for functional *in vivo* gene validation in the injected generation, we targeted the region orthologous to R141C in medaka embryos using the CRISPR/Cas9 system (Stemmer et al. 2015, Thumberger et al. 2021). As a negative control, we targeted the *oculocutaneous albinism* 2 locus (*oca2*) (Fukamachi et al. 2004) as a heart-unrelated pigmentation gene. The apparent loss of the eye pigmentation phenotype (and low degree of mosaicism) (Lischik et al. 2019; Hammouda et al. 2019) underscores the bi-allelic knock-out efficacy of our system. Injections into wild-type medaka embryos were performed at the 1-cell stage, and hereafter resulting embryos are referred to as crispants. To address the impact on the heart rate we raised the medaka crispants until cardiac function was fully developed and the heart rate had reached a plateau at 4 days post fertilization (Gierten et al. 2020) (4 dpf; developmental stage ~31-32 (Iwamatsu 2004)).

**Figure 1.**
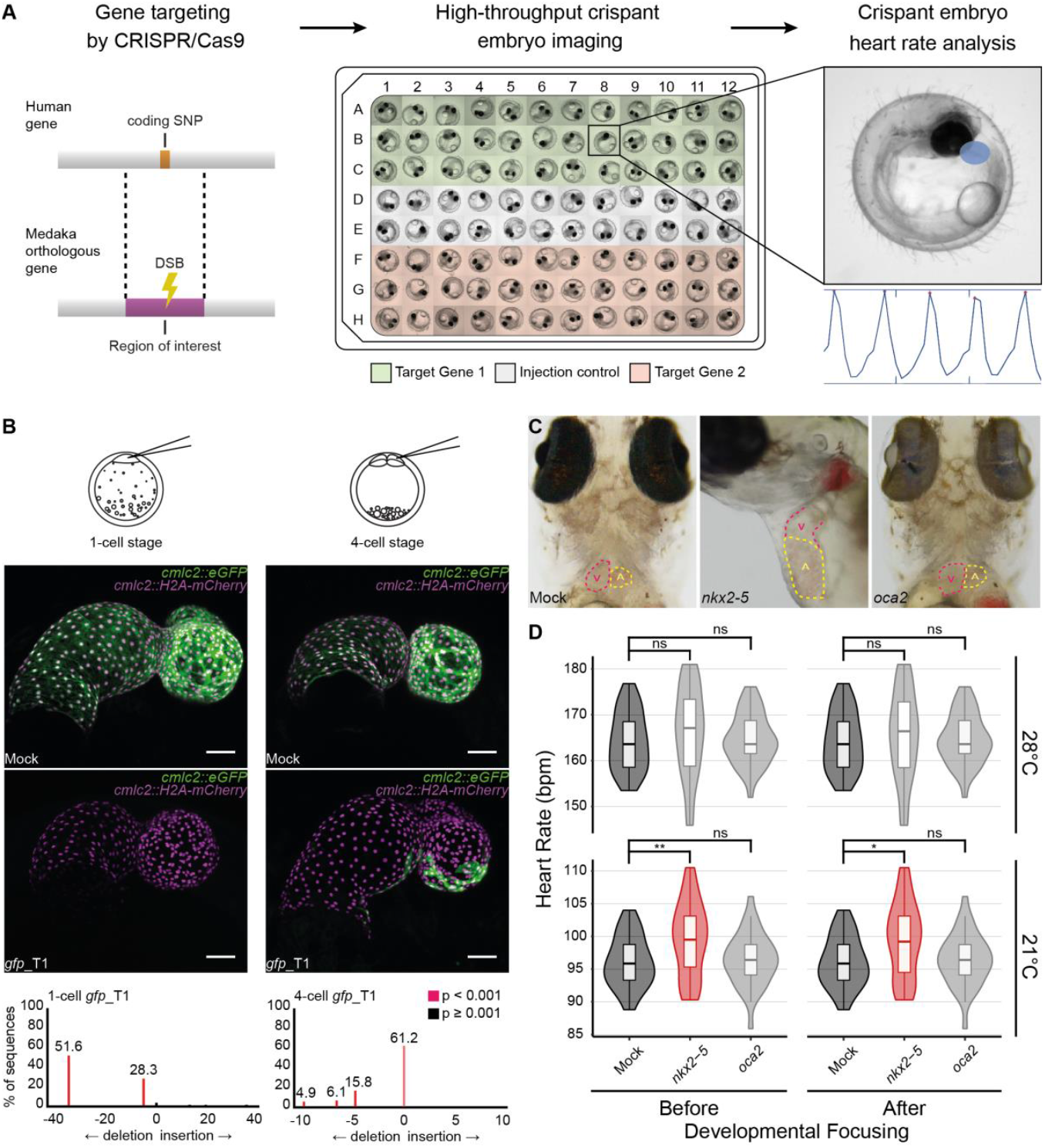
Functional gene validation pipeline confirms heart rate phenotype in *nkx2-5* crispants. **A** Schematic overview of our functional gene validation pipeline: position of human coding SNP mapped to medaka orthologous gene to define region of interest for CRISPR/Cas9 gene targeting (double strand break; DSB). 96-well plate layout of crispant embryos (Target Gene 1 and 2) separated by *GFP* mRNA mock-injected siblings. Embryos are subjected to high-throughput imaging followed by automated heart detection (blue area) and heart rate quantification (graphical output; *HeartBeat* software (Gierten et al. 2020). **B** Confocal images of GFP expression in mock-injected and *gfp* crispant embryo hearts of the *cmlc2::eGFP cmlc2::H2A-mCherry* reporter line (7 dpf). Embryos were injected either at the 1-cell or 4-cell stage. Note: complete loss of GFP expression when injected at the 1-cell stage (n = 8/8), while mosaic expression when injected at 4-cell stage (n = 4/4). Genotyping of *gfp* crispants display the genetic mosaicism resulting from Cas9-based targeting. Scale bars: 50 μm. For full image refer to Fig S1B. **C** Comparison of the atrium (A, dotted red line) and ventricle (V, dotted yellow line) in *GFP*-injected (Mock) and *nkx2-5* and *oca2* crispant embryos (9 dpf). Note: *nkx2-5* crispant shows dilated heart chambers while mock injected and *oca2* crispant embryos are indistinguishable. Loss of eye pigmentation in *oca2* crispants reflects high efficiency of knock-out rate in the injected generation. **D** Heart rate measurements (beats per minute, bpm) of *GFP*-injected (Mock; dark grey), *nkx2-5* and *oca2* crispant embryos (4 dpf) at 21 and 28°C, before (left) and after (right) exclusion of severely affected embryos (developmental focusing) reveal elevation of mean heart rates in *nkx2-5* targeted embryos, significant at 21°C (red). Significance was determined by two-tailed Student’s t-test; *\*p* < 0.05, *\*\*p* < 0.01, ns (not significant; light grey). For biological replicates see Source Data Fig 1D.

To assess changes in mean heart rate with statistical significance, we took advantage of a 96-well plate format, and imaged multiple biological replicates of crispants (3 rows; n=36 per condition) as well as of *GFP mRNA* mock-injected siblings as internal plate control (2 rows; n=24) (Fig S2A). To assess heart function under different environmental conditions, embryos were imaged at two different temperatures (21 and 28°C, respectively). Heart rates of all embryos were quantitatively determined from the imaging data using the *HeartBeat* software (Gierten et al. 2020), and randomly selected embryos were genotyped to correlate CRISPR/Cas9 targeting (Hammouda et al. 2019).

While mock injected embryos did not show phenotypes, crispants of the positive control *nkx2-5* displayed a variety thereof. These ranged from global severe developmental delays to local cardiac malformations morphologically resembling the phenotypes previously observed in zebrafish *nkx2-5* mutants such as enlarged heart chambers (Fig 1C) (Targoff et al. 2013). Notably, in the negative control (*oca2* crispants) neither cardiac nor developmental phenotypes were observed (Fig 1C), indicating that genome targeting of *oca2* as well as injection and handling of the embryos did not impact on heart and general development per se. Quantitative comparison of cardiac function revealed an overall elevation in the mean heart rate of *nkx2-5* crispants (21°C 99.7 bpm, 28°C 166 bpm) compared to mock control siblings (21°C 96.1 bpm, 28°C 164 bpm) with a significant (p = 0.0074) difference at 21°C (Fig 1D; left panel). Notably, independent experimental replicates targeting the same *nkx2-5* exon with two different sgRNAs robustly yielded a significant heart rate phenotype at 21°C (Fig S2B-C). In contrast, the mean heart rate in *oca2* crispants was indifferent from mock control at either temperature (21°C 96.4 bpm, 28°C 165 bpm), validating *oca2* as bona fide negative control.

To avoid severe developmental delays in *nkx2-5* crispants to potentially skew heart rate comparisons, we further applied a developmental focusing filter. Only embryos having developed beyond stage 28 (Iwamatsu 2004), at which cardiac function was previously shown to have reached a functional plateau (Gierten et al. 2020), were chosen for statistical analysis. Developmental focusing, only excluded three embryos from the *nkx2-5* group which did not impact on the results (Fig 1D; right panel). These results underline the robustness of our pipeline and demonstrate its sensitivity to detect mild heart rate phenotypes reflecting cardiac function already at embryonic stages of medaka development.

Next, we determined the baseline probability of heart rate phenotypes by targeting a set of randomly selected genes with CRISPR/Cas9. From a total of 23622 annotated medaka coding genes in Ensembl (Yates et al. 2020), we used a random number generator to select 10 genes (Table 1). For each gene, a random exon was chosen for targeting via CRISPR/Cas9. Heart rates of 36 crispants were assessed per locus. To control for potential heart rate fluctuations in embryos within and across different experiments (i.e. 96-well plates), we included mock-injected siblings as internal plate control. Heart rates of target gene crispants and control siblings were scored and the means were compared before and after developmental focusing.

**Table 1.**
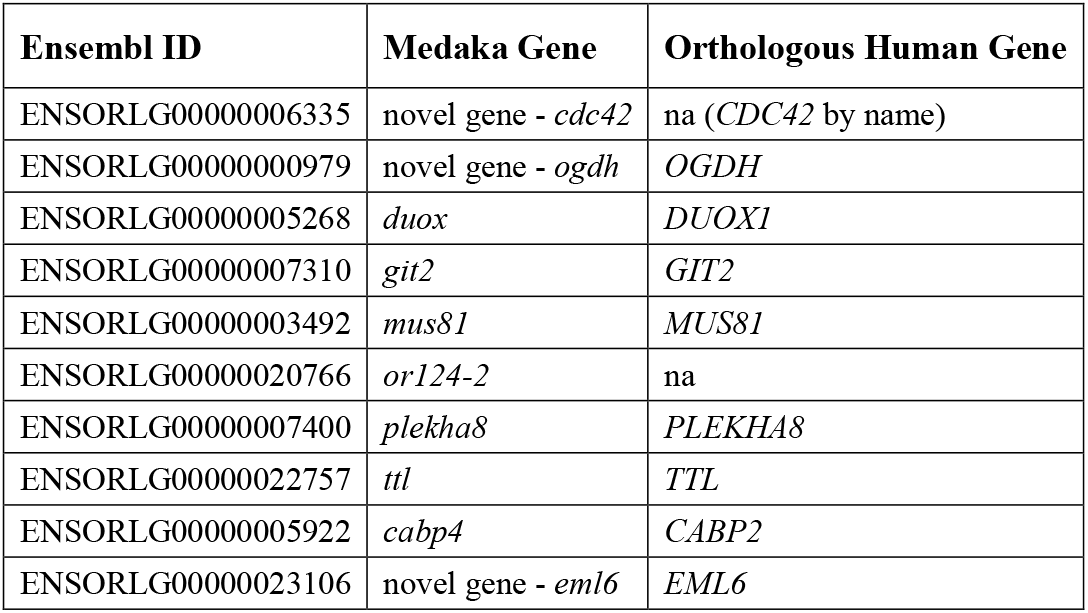
List of randomly selected genes. Medaka Ensembl gene names and codes, as well as orthologous human genes as annotated in the 95^th^ Ensembl release.

Comparative heart rate analysis revealed a heart rate phenotype in two out of the ten randomly selected genes at both temperatures measured (Fig 2, and Fig S3). Remarkably, both genes of the random set, the *oxoglutarate dehydrogenase (ogdh)* and the *cell division control protein 42 homolog (cdc42)* have been previously associated with heart phenotypes in human GWAS or were shown to play a role in heart function, respectively (Thanassoulis et al. 2013; Qian et al. 2011; Li et al. 2017). These results confirm the reliability of our assay to identify genes of a given set that affect cardiac function. Of note, repeated rounds of random gene selection revealed a similar mean baseline of 10-20% of selected genes being related to the heart in GWAS or previous experimental reports.

**Figure 2.**
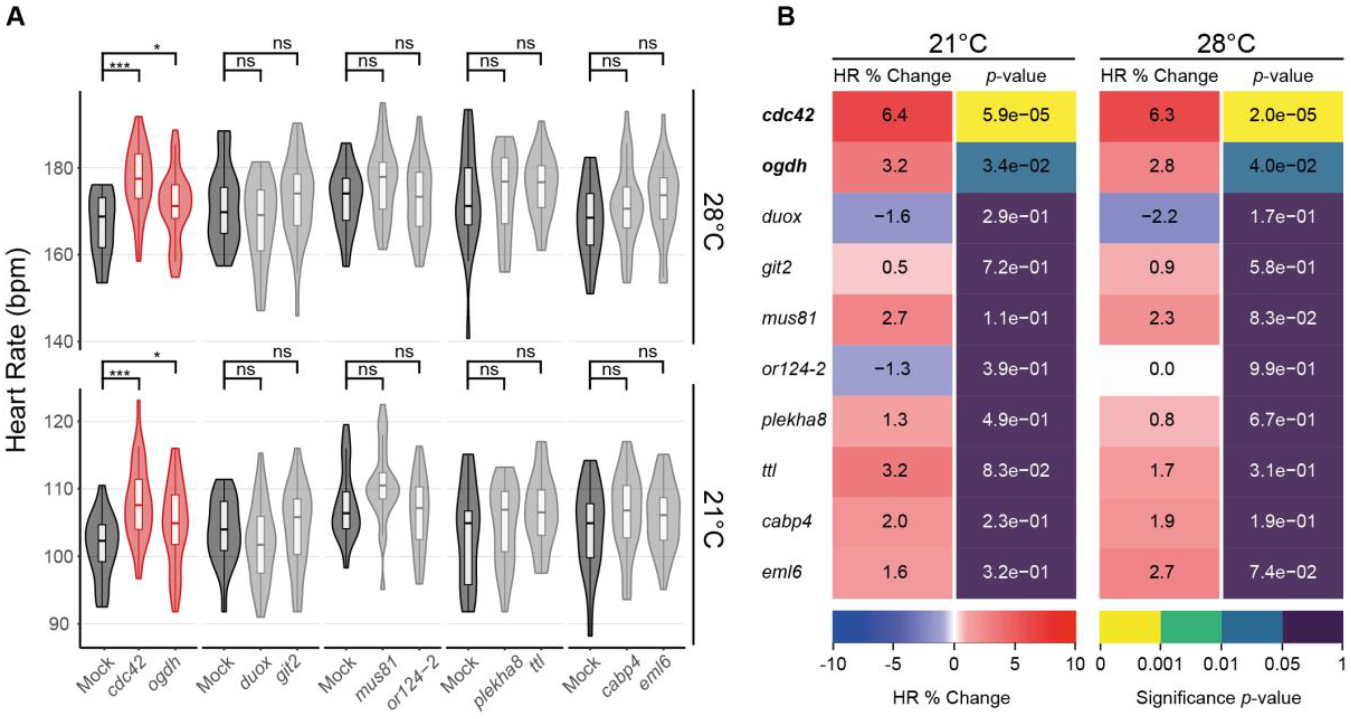
Baseline probability of heart rate phenotype assessed via *in vivo* targeting of randomly selected genes. **A** Heart rate measurements (beats per minute, bpm) of *GFP*-injected (Mock) and corresponding sibling crispant embryos (4 dpf) at 21 and 28°C after developmental focusing. Different experimental plates are represented by breaks on the x-axis. Significant differences in mean heart rates were determined between each crispant embryo group and its corresponding sibling control group by two-tailed Student’s t-test; \**p* < 0.05, \*\*\**p* < 0.001, ns (not significant). Crispants showing significant heart rate phenotype (red), *GFP*-injected controls (Mock; dark grey), crispants showing no significant heart rate phenotype (light grey). **B** Heatmap quantitative representation of the data shown in (A); for each measured temperature, the percent change in mean heart rate (HR % Change) between crispants and their corresponding control sibling, flanked by the statistical significance (*p*-value) of the observed change calculated by two-tailed Student’s t-test on the full distribution in (A). Genes showing significant heart rate phenotypes are indicated in **bold**. For biological replicates see Source Data Fig S3.

We next applied our pipeline to interrogate a larger, targeted selection of genes associated with cardiovascular diseases in human GWAS. We used GRASP (Eicher et al. 2015), the genomewide repository of associations between single nucleotide polymorphisms (SNPs) and phenotypes, to compile a list of 40 candidate genes from human GWAS with a coding association to heart phenotypes (hGWAS genes; Table 2). We focused on genes with no prior experimental link to heart function, while including few known heart genes as additional positive controls. To address the specificity of our approach, the selected candidate genes were categorized according to their association into general heart related (n=17) or related specifically to heart rate (n= 23; Table 2).

**Table 2.**
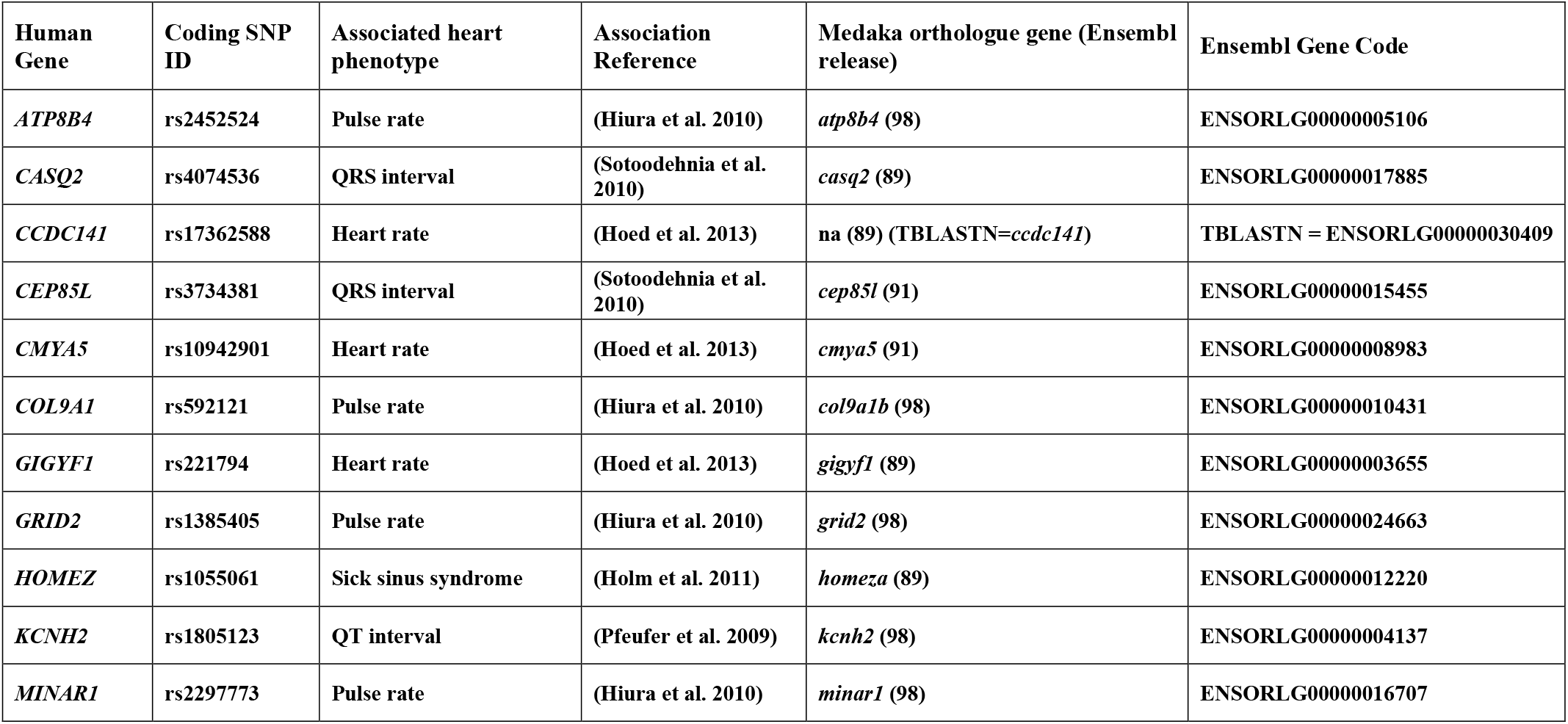
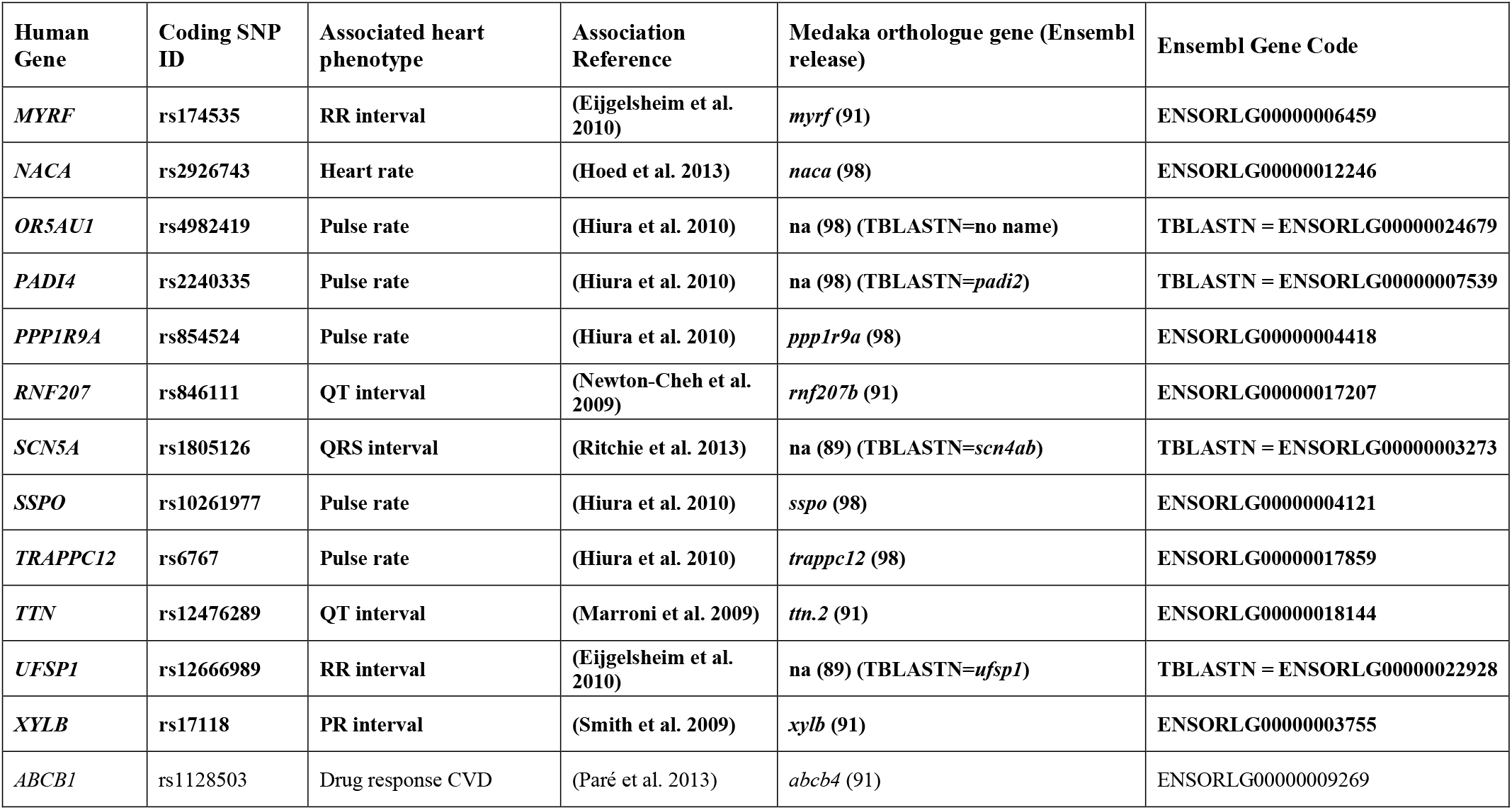
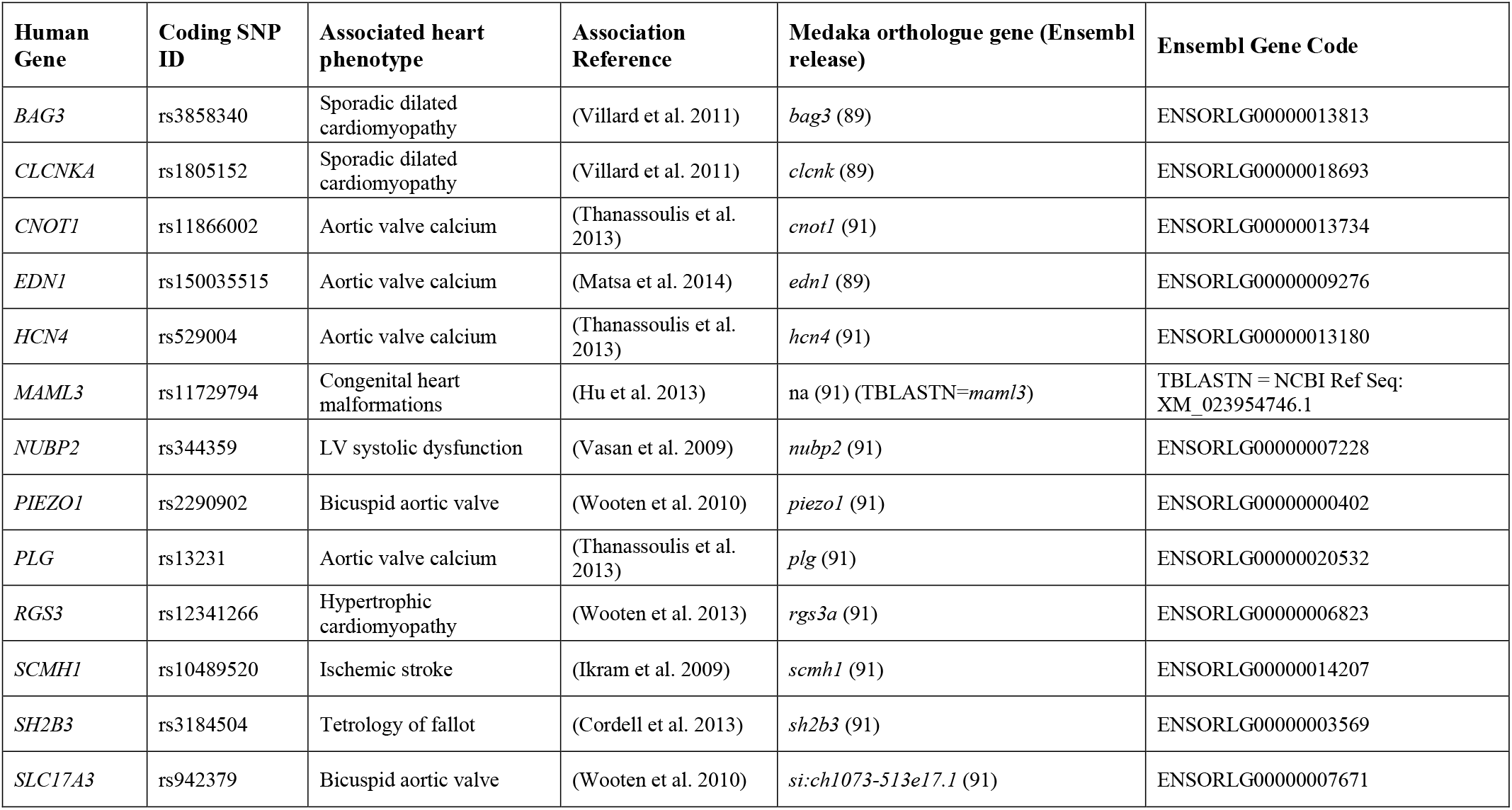
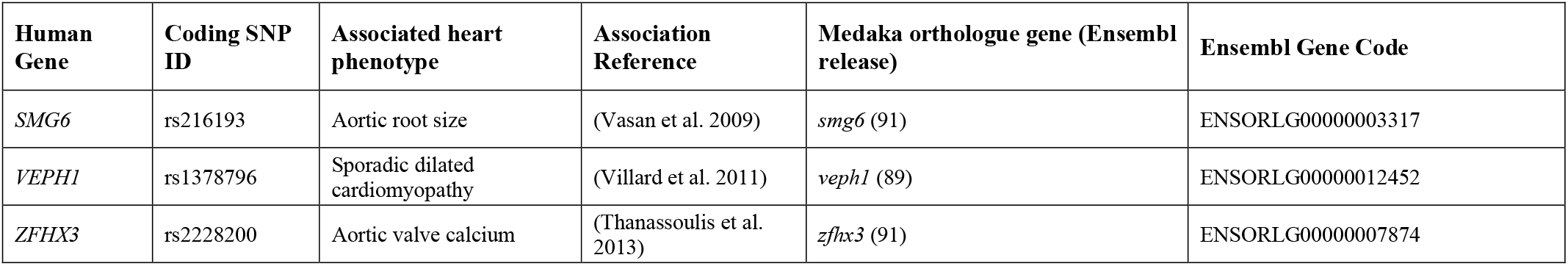
List of candidate genes extracted from Human GWAS using GRASP 2.0 Database. Human genes are categorized according to their association into “heart rate” (**bold**) and “non-heart rate” (non-bold) related phenotypes in human GWAS.

Heart rates of candidate gene crispants and control injected siblings were scored and compared before (Fig S5) and after developmental focusing (Fig 3A and Figure S4). Across the hGWAS set of 40 genes, comparative heart rate analysis showed statistically significant heart rate phenotypes in a total of 16 genes (Fig 3B). The five positive controls, known to play key roles in heart functions such as cardiac contraction *(TTN* (Itoh-Satoh et al. 2002) and *NACA* (Park et al. 2010)) and heart rate regulation *(CASQ2* (Faggioni & Knollmann 2012), *KCNH2* (Gianulis & Trudeau 2011) and *SCN5A* (Zaklyazminskaya & Dzemeshkevich 2016; Huttner et al. 2013)) clearly responded in the assay. Beyond known cardiac genes, we revealed new genes linked to various biological functions *(CCDC141, GIGYF1, HOMEZ, MYRF, SMG6, CMYA5, CNOT1, SLC17A3, TRAPPC12, SSPO* and *PADI4)* which up to now, had little to no experimental evidence in cardiac function (Rathjens et al. 2020; Benson et al. 2017; Yamaguchi et al. 2018; Elmen et al. 2020).

**Figure 3.**
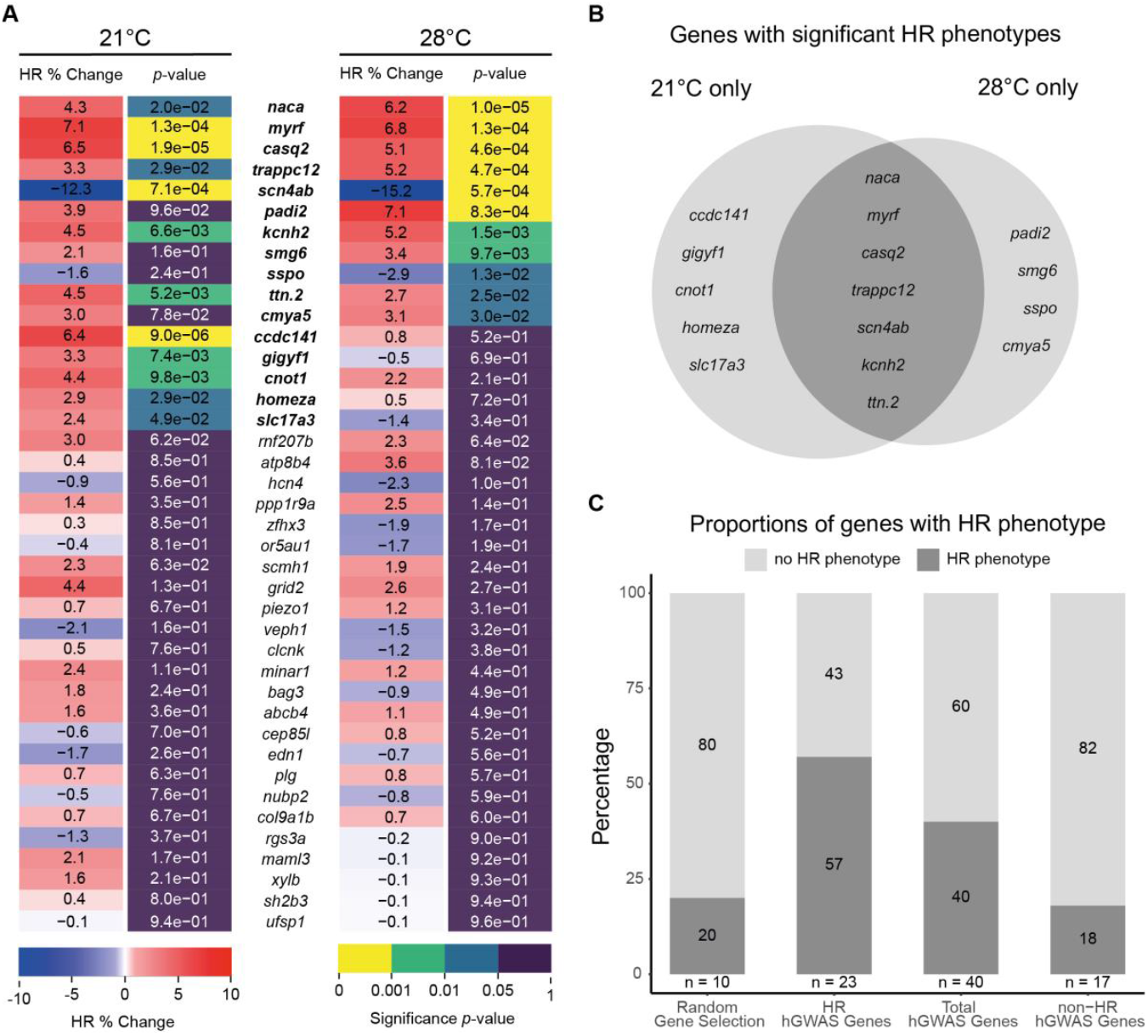
Targeted human heart-GWAS validations reveal new genes affecting heart rate. **A** Heatmap quantitative representation of the comparative heart rate analysis between each crispant embryo group and its corresponding control sibling group after developmental focusing (also see plots in Fig S4); for each measured temperature, the percent change in mean heart rate (HR % Change) between crispants and their corresponding control sibling, flanked by the statistical significance (*p*-value) of the observed change calculated by two-tailed Student’s t-test on the full distribution (Fig S4). Genes showing significant heart rate phenotypes are indicated in **bold**. For biological replicates see Source Data Fig S4. **B** Venn diagram summarizing the genes with significantly different heart rate (HR) phenotypes only at 21°C, only at 28°C or at both temperatures (dark grey). **C** Stacked plots representing percentage of genes showing a significant heart rate phenotype (dark grey) in each group. Number of genes for each group is denoted (n). hGWAS corresponds to the selection of genes associated to heart phenotypes in human GWAS.

When analyzing the candidates according to their GWAS association (“heart rate” and “non – heart rate” phenotypes), we observed a strong positive correlation between the respective phenotypes observed in medaka crispants already at the embryonic stages used and the associated phenotype in adult human GWAS. The proportion of heart rate-associated genes in hGWAS that yield a heart rate phenotype in medaka embryos (13/23) was elevated compared to the proportion of non-heart rate-associated genes yielding a heart rate phenotype (3/17). Even when considering the entire group, we observed a higher proportion of genes with an effect on heart rate in our targeted hGWAS gene set (16/40) compared to our randomly selected gene set (2/10) (Fig 3C). Taken together, phenotypes in early medaka embryos likely reflect risk factors in human adults, thus we uncovered functionally relevant heart rate phenotypes in previously uncharacterized genes.

In addition to the observed heart rate phenotypes in *trappc12* crispants, we also uncovered morphological heart phenotypes such as heart looping defects. Where in wild-type medaka, heart looping usually starts at around stage 27, when the atrium shifts to the right and lies adjacent to the ventricle (Iwamatsu 2004), *trappc12* crispants revealed strong heart looping phenotypes (12/46) not observed in *oca2* crispants (0/22) or mock-injected embryos (0/26) (Fig 4A).

**Figure 4.**
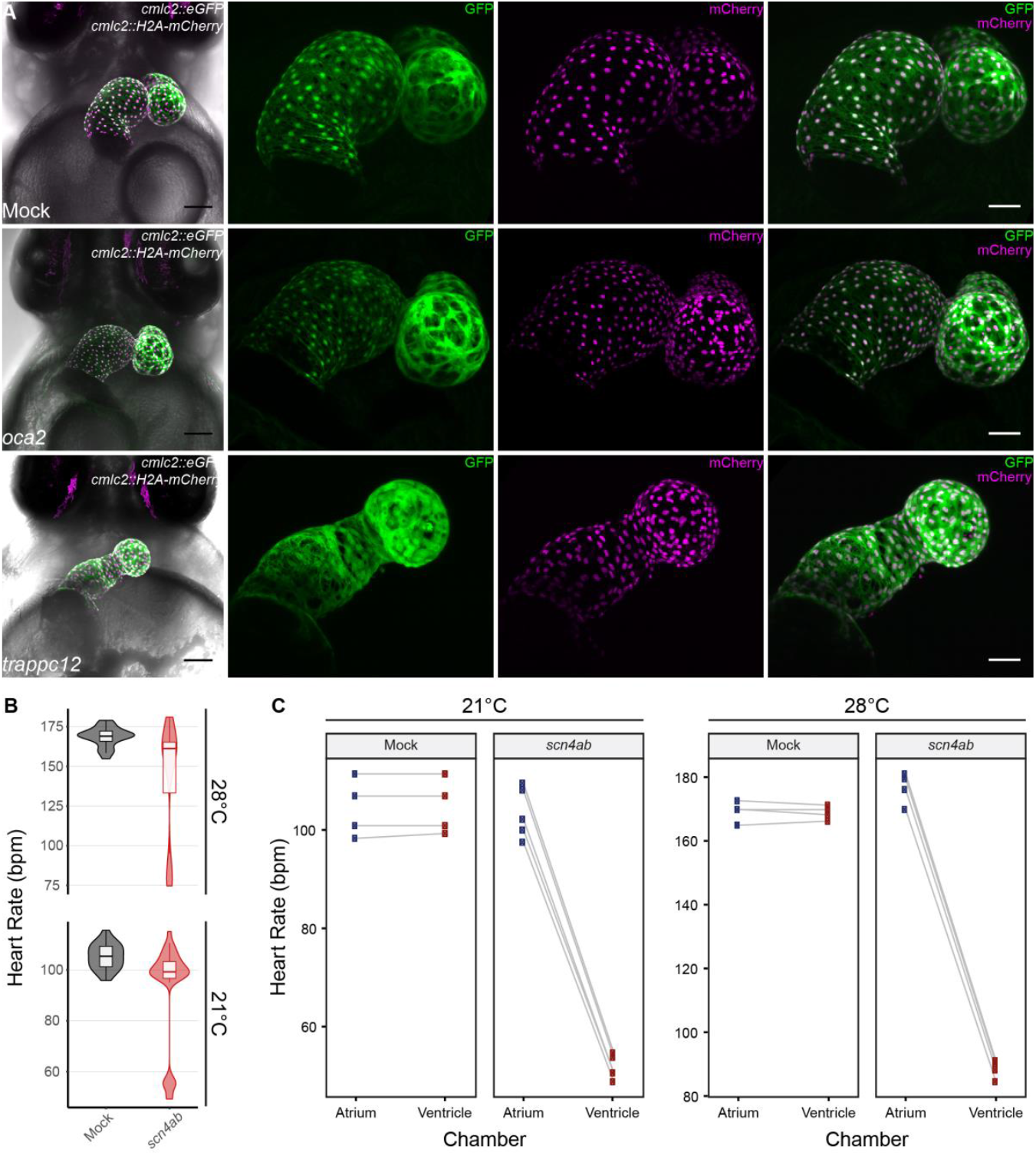
Heart looping defects and AV-block type arrhythmia in medaka crispant embryos. **A** Confocal images of hearts of mock-injected, *oca2* and *trappc12* crispants of the *cmlc2::eGFP cmlc2::H2A-mCherry* reporter line (7 dpf); note the heart looping defect observed in *trappc12* crispants. Scale bars: 100 μm (First panel on left) and 50 μm (blow-up images). **B** Heart rate measurements (beats per minute, bpm) of *GFP*-injected (Mock; dark grey; n = 22) and *scn4ab* crispant (red; n = 32) embryos (4 dpf) at 21 and 28°C; note the bimodal distribution in *scn4ab* crispants. **C** Paired plots showing heart rate scores for each chamber separately (atrium in blue; ventricle in red) of individual embryos from B at both temperatures.

In crispants of *scn4ab,* heart rate analysis interestingly uncovered a bimodal distribution, with a population displaying roughly half the average heart rate at both recorded temperatures (Fig 4B). Visual inspection of the *scn4ab* crispant embryos revealed an arrhythmic heart anomaly previously reported in zebrafish *scn5a* mutants (Huttner et al. 2013), i.e. known as an atrioventricular block (AV-block), characterized by a delay or disruption of impulse transmission from the atrium to the ventricle. Scoring the beat frequency of both heart chambers separately in individual embryos exposed the impaired rhythm of atrial to ventricular contractions, which resulted in a delay or even skipping of ventricular beats in the *scn4ab* crispants but not in control siblings (Fig 4C). *scn4ab* crispants displayed various severities of the AV-block from mild (regular heart beats with occasional beat skipping), to moderate (consistent 2:1 atrial to ventricular contraction; Movie S1), to severe (3:1 or more; Movie S2). Still heavily affected *scn4ab* crispants survived until hatching. Impressively, the prevalence of the arrhythmia phenotype in *scn4ab* crispants was markedly high, exceeding 90% of the injected embryos, reflecting the high efficiency of the *heiCas9* and the high penetrance of the mutations introduced. Notably, the arrhythmia phenotype of *scn4ab* mutants bred to homozygosity did not differ from the phenotype observed in the injected generation F0 (Movie S1), verifying the specificity of the phenotype. These results further underscore the efficacy and reliability of medaka F0 crispant analysis as a rapid validation tool to identify genes with a functional link to human cardiac diseases.

## Discussion

Most cardiovascular diseases can be prevented if diagnosed and treated early. Previous studies have shown the importance of the resting heart rate as a vital risk factor both in terms of prediction and prevention of CVDs (DYER et al. 1980; MD et al. 2010; Eppinga et al. 2016). An increase of 5 beats per minute correlates with a 20% increase in risk of mortality (Eppinga et al. 2016), and reducing the resting heart rate has proven to improve the clinical outcomes of various CVDs (Beere et al. 1984; MD et al. 2010).

Human GWAS have been performed in search of genetic determinants of CVDs, and although a wide array of candidate genes with various functions are being associated to heart phenotypes in human GWAS, further focus is usually turned to those few genes with pre-existing indication of cardiac function. Other associated genes with an unknown function or without a pre-existing functional link to the heart are often neglected and not pursued further, pushing all those into a blind spot, resulting in a negative loop of discovery. Thus, it is important to address the role of such genes in heart function through experimental validation in model organisms, in pursuit of novel markers for CVD diagnosis. We address this blind spot of discovery by applying a high-throughput heart rate imaging and analysis pipeline coupled to a reverse genetic validation approach via CRISPR/Cas9 mediated mutagenesis in a genetically suited vertebrate model (Fig 1A).

F0 mutagenesis screens are becoming more and more popular (Teboul 2017; Shankaran et al. 2018; Wu et al. 2018; Heyde et al. 2020), largely due to the improvements in CRISPR/Cas9 gene targeting efficiency by modification of the enzyme or promoting nuclear localization (Thumberger et al. 2021). Overall, gene targeting in medaka using CRISPR/Cas9 has proven to be highly efficient, as shown by the prominent loss of GFP expression in *gfp* crispants when injected at the 1-cell stage (Fig 1B and Fig S1) and the prominent bi-allelic mutagenesis as apparent by the loss of eye-pigmentation in the *oca2* crispants (Fig 1C and Fig S2A). A similarly high penetrance was also observed in the *scn4ab* crispants, where we detected and quantified severe arrhythmia phenotypes such as AV-block with our assay (Fig 4B-C, Movies S1 and S2). A 90 % prevalence of the arrhythmia phenotype and an absence of global phenotypes further reflect the specificity of this phenotype.

A subset of the target genes in our assay (e.g. *nkx2-5, smg6, naca, ttn.2, abcb4)* however, yielded a rather broad range of global developmental phenotypes, potentially reflecting their important roles in embryonic development. To address this un-avoidable outcome when tackling genes with broader function (e.g. transcription factors or essential genes), we applied a developmental focusing filter in the analysis phase. Doing so, we avoid a biased assessment of the heart rate by ensuring the comparability of the crispant embryos on a global developmental scale, which in turn allows emphasizing cardiac-specific effects. Interestingly however, developmental focusing, although deemed important, in only few cases significantly altered the outcome of the analysis (Fig. S5). This reflects the robustness of the assay and the homogeneity of CRISPR/Cas9-induced phenotypes in the isogenic background of the medaka line used.

In a set of ten randomly chosen genes we observed a baseline occurrence of heart-affecting genes of about 20 % (Fig 2, and Fig S3). Relevantly, both genes were implicated in heart functions, further reflecting the reliability of our model and approach. For *cdc42,* there is *a priori* evidence of its human orthologue in heart development as well as in regulating heart function across species (Qian et al. 2011; Li et al. 2017). Surprisingly, we did not find any associations (coding or non-coding) of *CDC42* to heart phenotypes in human GWAS according to the GRASP database (Eicher et al. 2015). As for *ogdh,* no experimental evidence in cardiac function has been previously reported, but a polymorphism located on one of its exons has been associated to heart phenotypes in human GWAS (Thanassoulis et al. 2013). Except for *duox,* which has been reported as having an indirect role in cardiac regeneration in zebrafish (Han et al. 2014), none of the other randomly selected genes were, to our knowledge, ever connected with cardiac function. In summary, from our random gene set, the only two affecting the heart beat (*ogdh* and *cdc42*) are connected to cardiac function by prior evidence (Thanassoulis et al. 2013; Qian et al. 2011; Li et al. 2017).

All but one positive (*HCN4*) control among the hGWAS candidates resulted in a pronounced heart rate phenotype in our assay, reflecting their role in cardiac contraction *(TTN* (Itoh-Satoh et al. 2002) and *NACA* (Park et al. 2010)), cardiac conduction and heart rate regulation (*CASQ2* (Faggioni & Knollmann 2012), *KCNH2* (Gianulis & Trudeau 2011) and *SCN5A* (Zaklyazminskaya & Dzemeshkevich 2016; Huttner et al. 2013)). In case of the missing heart rate phenotypes anticipated in *hcn4* crispants, we suspect compensation by its paralog *hcn4l*. For eleven hGWAS candidate genes, our analysis provided the first experimental evidence validating a cardiac function, and accordingly put these identified genes under the spotlight as new targets for future in-depth characterization and as candidates for the prediction of heart diseases prior to their onset. This is impressively substantiated by the emerging studies on the CCR4-NOT *(CNOT1)* complex in heart structure and function (Yamaguchi et al. 2018; Elmen et al. 2020).

Throughout our study, we primarily focused on the heart rate as a measure of cardiac function due to its ease of quantification and interpretation in high-throughput. However, we did also notice heart morphological phenotypes in some crispants across our hGWAS candidates list. Among others, *trappc12* crispants showed heart looping defects, resulting in the improper placement of the atrium compared to the ventricle (Fig 4A). With advances in AI-based image analysis, important morphological features could be additionally extracted in high throughput, leading to a better understanding of factors involved in heart development and function. Grouping the candidate genes according to their heart GWAS association into heart rate and non-heart rate-related phenotypes further exposed the prominent positive correlation between the associated human phenotype and the observed phenotype in medaka (Fig 3C). Medaka’s isogenic background, a product of inbreeding over multiple generations (Wittbrodt et al. 2002), enabled the detection of subtle changes in heart rate immediately in F0 crispants. This accelerated the analysis and avoided the necessity to analyze homozygous offspring in the second and third generation after CRISPR targeting.

It is noteworthy that despite the evolutionary distance from fish to humans the medaka phenotypes match the class of hGWAS effects. This is even more relevant since the roles of the genes in medaka were validated in embryos, suggesting that the validated marker genes have predictive power in humans. This deep functional conservation emphasizes the potential of our approach for the identification and validation of novel predictive genetic markers for cardiovascular diseases in humans. We have showcased a highly versatile, sensitive and robust high-throughput reverse genetic validation assay to address the pool of understudied, neglected putative candidates. In the future, the combination of genetic validation and drug screening in a single platform building on our assay will facilitate the simultaneous identification of novel genetic players and interacting small molecules with rescuing power.

## Materials and Methods

### Ethics Statement

All fish are maintained in closed stocks at Heidelberg University. Medaka (*Oryzias latipes*) husbandry (permit number 35–9185.64/BH Wittbrodt, Regierungspräsidium Karlsruhe) was performed according to local animal welfare standards (Tierschutzgesetz §11, Abs. 1, Nr. 1) in accordance with European Union animal welfare guidelines (Bert et al. 2016). The fish facility is under the supervision of the local representative of the animal welfare agency. The following medaka stocks and transgenic lines were used: wild-type Cabs and *cmlc2::eGFP cmlc2::H2A-mCherry* transgenic HdrR-II strain. Medaka embryos were used at stages prior to stage 42. Medaka were raised and maintained as described previously (Köster et al. 1997).

### Generation of the transgenic cmlc2 dual-reporter line

For dual-color cardiac imaging, *cmlc2::eGFP* and *cmlc2::H2A-mCherry* transgenic medaka lines were generated in the wild-type HdrR-II background. A modified version of pDestTol2CG (http://tol2kit.genetics.utah.edu/index.php/PDestTol2CG) was used containing a *cmlc2::eGFP* reporter cassette. For the second, nuclear reporter, the *eGFP* was replaced by an *H2A-mCherry* insert. Both plasmids were each co-injected at 10 ng/μl with 10 ng/μl Tol2 transposase mRNA into HdrR-II one-cell stage embryos using the microinjection technique as previously described (Rembold et al. 2006) to generate separate reporter lines. A double transgenic line was derived from a cross of the *cmlc2::eGFP* line to *cmlc2::H2A-mCherry* line and maintained for CRISPR-Cas9 injections.

### Candidate gene selection

For the unbiased gene targeting, an online random number generator was used to generate 10 numbers between 1 and 23622, corresponding to the number of annotated medaka coding genes in Ensembl (Yates et al. 2020) (Table 1). The number of exons for each gene was counted and a random number was generated to select the exon for CRISPR/Cas9 targeting. For the targeted human heart-GWAS (hGWAS) gene selection, the genome-wide repository of associations between SNPs and phenotypes (GRASP v2.0) was used (Eicher et al. 2015). In the search field, “Heart” and “Heart rate” were chosen as the respective categories for all heart- and heart rate-related phenotypes associated in human GWAS, only coding SNPs (i.e. SNP functional class = exons) were searched for. List of resulting genes was extracted (Table 2), and candidate genes for the functional validation assay were chosen. The focus was on uncharacterized genes, or genes with no prior experimental link to heart function, yet some known heart genes were included as proof of concept. For each hGWAS candidate gene, the corresponding medaka ortholog was extracted using Ensembl (Yates et al. 2020). For the few genes which did not have an annotated medaka ortholog, the human protein sequence was BLASTed using the “tblastn” function of the NCBI BLAST (https://blast.ncbi.nlm.nih.gov/Blast.cgi) and Ensemble (http://www.ensembl.org/Multi/Tools/Blast) online tools to obtain a target medaka locus. Using Geneious 8.1.9 (https://www.geneious.com), regions of interest (ROI) on medaka orthologous genes for CRISPR targeting were primarily chosen based on the corresponding location of human SNP when aligning the medaka and human protein sequences.

### sgRNA target sites selection and in vitro transcription

All sgRNA target sites used in this study are listed in Table S1. sgRNAs were designed with CCTop as described in Stemmer et al. (Stemmer et al. 2015). sgRNA target sites were selected based on number of potential off-target sites and their corresponding mismatches. Preferably, sgRNAs selected had no off-target site or at least 3 nucleotide mismatches. sgRNA for *oca2* was the same as in Lischik et al. (Lischik et al. 2019). Cloning of sgRNA templates and *in vitro* transcription was performed as detailed in Stemmer, et al. (Stemmer et al. 2015). All sgRNAs were initially tested after synthesis for *in vivo* targeting via injections into medaka embryos, followed by genotyping using our filter-in-tips protocol (Hammouda et al. 2019), in brief terms, by PCR amplification of target locus followed by T7 Endonuclease I assay (New England Biolabs).

### Microinjection

Medaka one-cell or four-cell stage embryos were injected into the cytoplasm as previously described (Stemmer et al. 2015). Injection solutions for CRISPR targeting comprised: 150 ng/μl heiCas9 mRNA (Thumberger et al. 2021), 15 ng/μl respective sgRNA and 10 ng/μl GFP mRNA as injection tracer. Control siblings were injected with 10 ng/μl GFP mRNA only. Injected embryos were incubated at 28 °C in embryo rearing medium (ERM), screened for GFP expression at 1 dpf and transferred to methylene blue-containing ERM (or plain ERM for reporter lines) and incubated at 28 °C until heart rate analysis (4 dpf) or confocal microscopy (7 dpf).

### Sample preparation and Imaging

For the heart rate assay, one day prior to imaging (3 dpf), medaka embryos were transferred from methylene blue-containing ERM into plain ERM and incubated at 28 °C. On day of imaging (4 dpf), individual medaka embryos (36 per sgRNA and 24 control injected) were administered to a 96 U-well microtiter plate (Nunc, Thermofisher #268152) containing 200 μl ERM per well and sealed using gas-permeable adhesive foil (4titude, Wotton, UK, 4ti-0516/96). Plates were automatically imaged using an ACQUIFER Imaging Machine (DITABIS AG, Pforzheim, Germany) at 21 and 28 °C with a 30-minute equilibration period before each measurement. Images were acquired in brightfield using 130 z-slices (dz = 0 μm) and a 2x Plan UW N.A. 0.06 objective (Nikon, Düsseldorf, Germany) to capture the centered embryo. Integration times were fixed with 80 % relative white LED intensity and 10 ms exposure time. Therefore, the whole 96-well plate was captured, with image sequences (videos) of entire microwells of approx. 10 seconds with 13 frames per second (fps). More details can be found in Gierten, et al. (Gierten et al. 2020).

For the live confocal microscopy of the reporter lines, injected embryos were treated from 4 dpf onwards with 5x phenylthiourea (PTU) in ERM solution to prevent pigmentation. On the day of imaging (7 dpf), PTU solution was washed away with ERM, embryos were rolled on fine sand paper and de-chorionated by incubation in hatching enzyme. Following de-chorionation, embryos were treated with 50 mM 2,3-butanedione monoxime (BDM) in 1x Tricaine solution until de-coupling of heart beat (~40 mins), which resulted in fully dilated heart chambers. Embryos were mounted on Matek dishes in 1.5% low-melting agarose with 85 mM BDM in 3x Tricaine solution. To avoid dehydration, mounted samples were covered with 30 mM BDM in 1x Tricaine solution throughout the imaging session. All confocal microscopy images were acquired at a Leica TCS SP8 with 10x dry or 20× oil objective, z-stacks of 200-300 μm were acquired with a z-step of 5 μm or 1 μm for 10x and 20x acquired images, respectively.

### *HeartBeat* detection and data analysis

Image optimizations prior to analysis, as well as heart rate analysis using the *HeartBeat* software were performed as previously described (Gierten et al. 2020). In some instances, heart rates could not be scored due to inconvenient embryo orientations shielding the view of the heart. For *scn4ab* crispants with cardiac arrhythmias, atrium and ventricle for individual embryos were separately segmented, and the respective beating frequency for each chamber was measured. Data plots were generated using ggplot2 package (Wickham 2016) in R 3.6.1 (R Core Team, 2019) and R-studio 1.2.1335 (RStudio Team, 2018). Statistical analysis for heart rate comparisons were computed in R. Significant differences were determined by two-tailed Student’s t-test. Significant *p*-values are indicated with asterisks (*) with \**p* < 0.05, \*\**p* < 0.01, \*\*\**p* < 0.001 and ns (not significant). Maximum intensity projections of confocal microscopy images were processed via Fiji image processing software.

### Embryo genotyping

Nucleic acid extraction and genotyping of embryos was done as previously described (Hammouda et al. 2019). Briefly, after imaging, embryos in 96-well plate were lysed in 50 μl Milli-Q water + 50 μl Fin-Clip lysis buffer each (0.4 M Tris-HCl pH 8.0, 5 mM EDTA pH 8.0, 0.15 M NaCl, 0.1 % SDS in Milli-Q water) using a custom 96-well mortar. The mortar was precleaned by incubation in hypochlorite solution (1:10 dilution of commercial bleach reagent) for at least 15 minutes followed by 5 minutes incubation in Milli-Q water. Plates containing lysed embryos were stored at 4 °C until genotyping. To confirm CRISPR on-target activity, per experimental plate, 2 embryos per condition were chosen at random for genotyping by PCR amplification of target locus using our filter-in-tips approach (Hammouda et al. 2019), followed by T7 Endonuclease I Assay (New England Biolabs). 30 PCR cycles were run in all samples, all primers used for PCR are listed in Table S2. Annealing temperatures were calculated using the online NEB Tm calculator (https://tmcalculator.neb.com/).

## Supporting information

Hammouda_Movie_S1

Hammouda_Movie_S2

## Acknowledgments

We thank A. Cornean for providing the *gfp_*T1 guide RNA and for his help in acquiring the confocal images. We thank V. Weinhardt, E. Tsingos and all members of the Wittbrodt lab for their critical, constructive feedback on the procedure and the manuscript, F. Loosli and S. Lemke for their constructive feedback towards the project as well as J. Backs and E. Furlong for an outside perspective. We thank T. Kellner for excellent technical support. We acknowledge the excellent fish husbandry of E. Leist, M. Majewsky and A. Saraceno. We thank J. Gehrig (ACQUIFER Imaging GmbH) for supporting us with the Imaging machine. O.T.H. is and J.G. was a member of the Heidelberg Biosciences International Graduate School (HBIGS). This work was supported by a fellowship of the Deutsches Zentrum für Herz-Kreislauf-Forschung (DZHK) to O.T.H and through DFG CRC-1324 TP B4 and NIH 5R01ES029917 – 03, KiyosuTox. J.G. was supported by the Deutsche Herzstiftung e.V. (S/02/17) and by an Add-On Fellowship for Interdisciplinary Science of the Joachim Herz Stiftung.

## Author contributions

O.T.H., T.T. and J.W. designed the study and implemented the methodology. J.G. generated the transgenic *cmlc2* dual-reporter line. O.T.H., V.K. and M.Y.W. synthesized the guide RNAs. O.T.H. performed experiments. O.T.H. analyzed the data with contributions by M.Y.W. and V.K.. O.T.H. visualized data. O.T.H. and J.W. wrote the first draft of the manuscript. O.T.H., T.T., and J.W. finalized the manuscript. J.W. provided resources and supervised this work.

---

**Figure S1.**
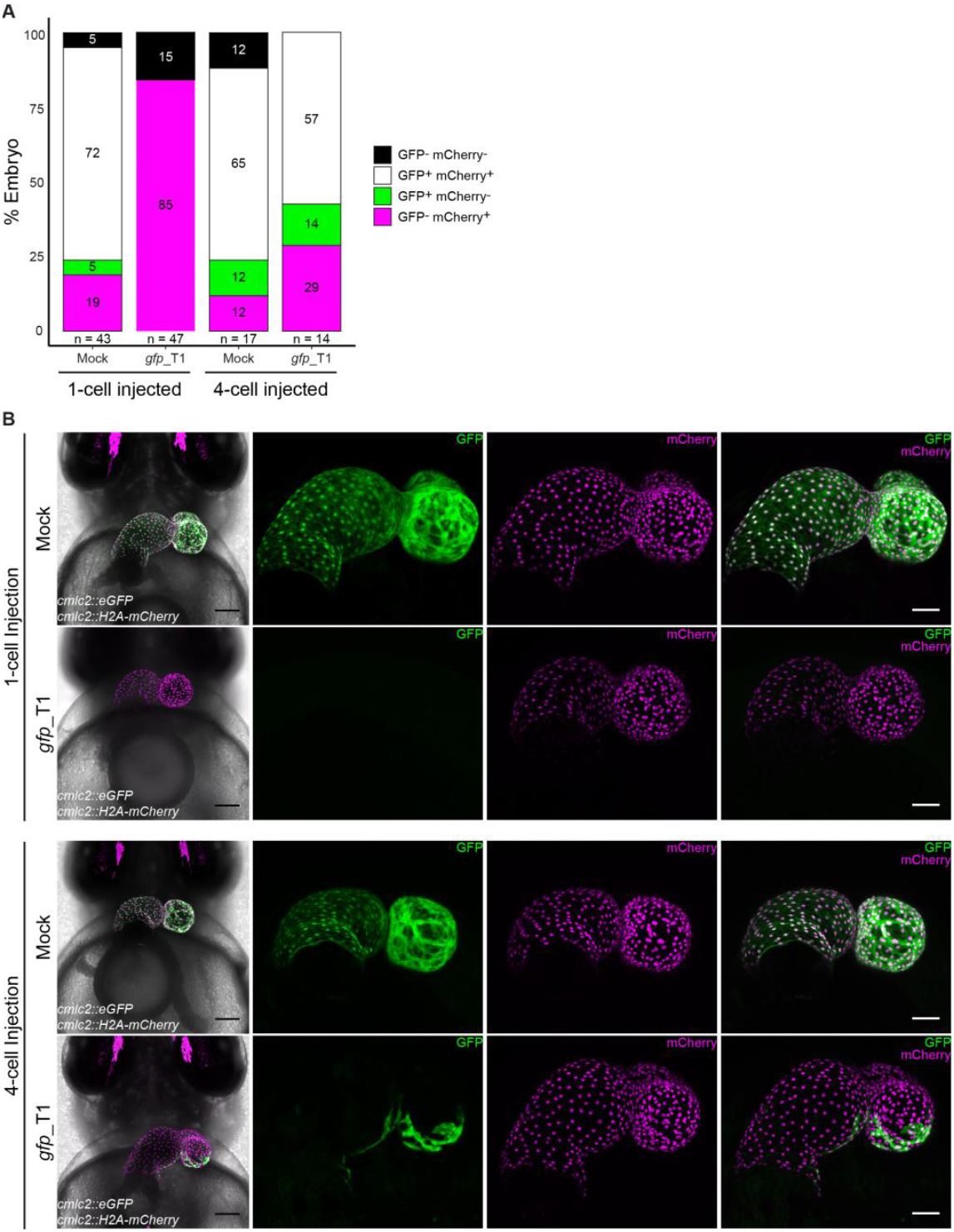
Highly efficient CRISPR/Cas9-mediated editing in injected (F0) generation. **A** Distribution of reporter expression in mock injected and *gfp_T1* crispants of the *cmlc2::eGFP cmlc2::H2A-mCherry* reporter line (4 dpf). Embryos were injected either at the 1-cell or 4-cell stage. Note: complete lack of GFP-expressing embryos when injected at the 1-cell stage. Biological replicates for each group is denoted (n). **B** Confocal images of GFP expression in mock-injected and *gfp* crispant embryo hearts of the *cmlc2::eGFP cmlc2::H2A-mCherry* reporter line (7 dpf). Embryos were injected either at the 1-cell or 4-cell stage. Note: complete loss of GFP expression when injected at the 1-cell stage (n = 8/8), while mosaic expression when injected at 4-cell stage (n = 4/4). Scale bars: 100 μm (First panel on left) and 50 μm (blow-up images).

**Figure S2.**
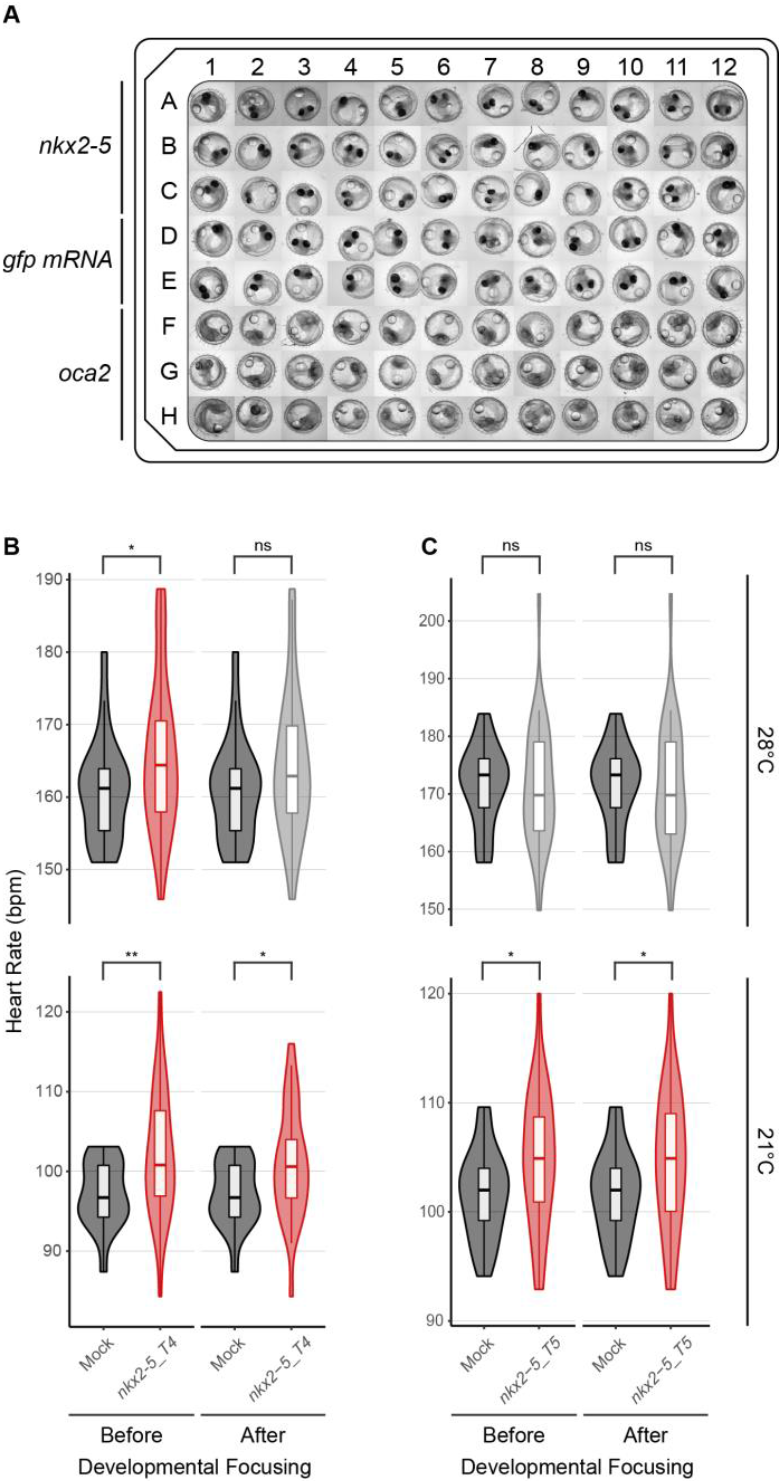
Consistent heart rate phenotype observed in medaka *nkx2-5* crispants. **A** Overview of 96-well plate with embryos (4 dpf) injected with sgRNA against *nkx2-5* or *oca2,* as well as embryos mock injected with *GFP mRNA* (Fig 1D). Note the loss of eye pigmentation in *oca2* crispant embryos. **B-C** Heart rate measurements of *GFP*-injected (Mock; dark grey) and *nkx2-5* crispant embryos (4 dpf) (**B** second replicate of *nxk2-5_T4*; **C** different sgRNA *nkx2-5_T5* targeting same region of interest) at 21 and 28°C, before and after exclusion of severely affected embryos (< stage 28; developmental focusing). Significant differences are shown in red and were determined by two-tailed Student’s t-test; \**p* < 0.05, \*\**p* < 0.01, ns (not significant; light grey). For biological replicates see Source Data Fig S2.

**Figure S3.**
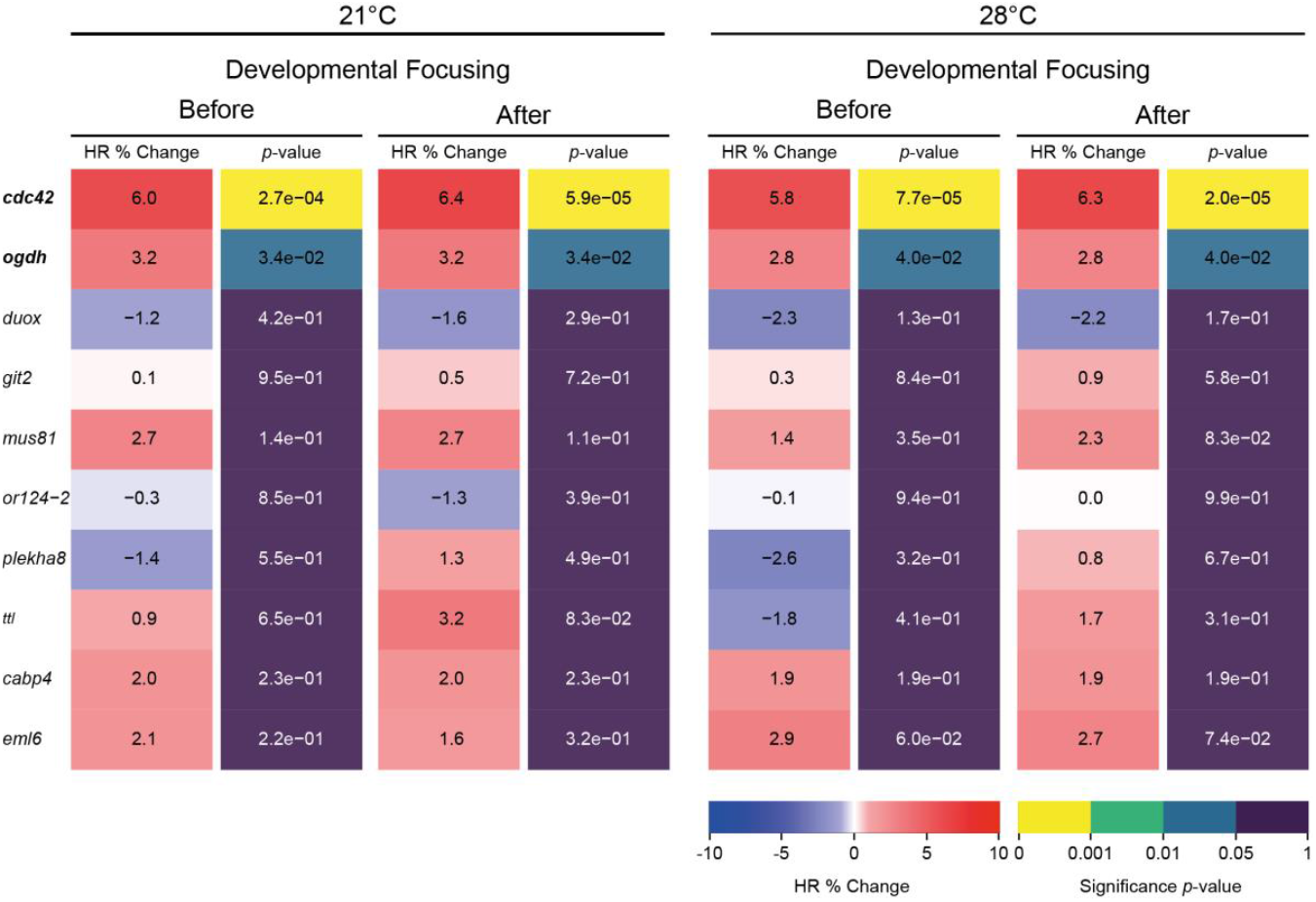
Developmental focusing does not alter analysis outcome of random gene selection. Heatmap quantitative representation of the comparative heart rate analysis between each crispant embryo group and its corresponding control sibling group before and after developmental focusing; for each measured temperature, the percent change in mean heart rate (HR % Change) between crispants and their corresponding control sibling, flanked by the statistical significance (*p*-value) of the observed change calculated by two-tailed Student’s t-test on the full distribution. Genes showing significant heart rate phenotypes are indicated in **bold**. For biological replicates see Source Data Fig S3.

**Figure S4.**
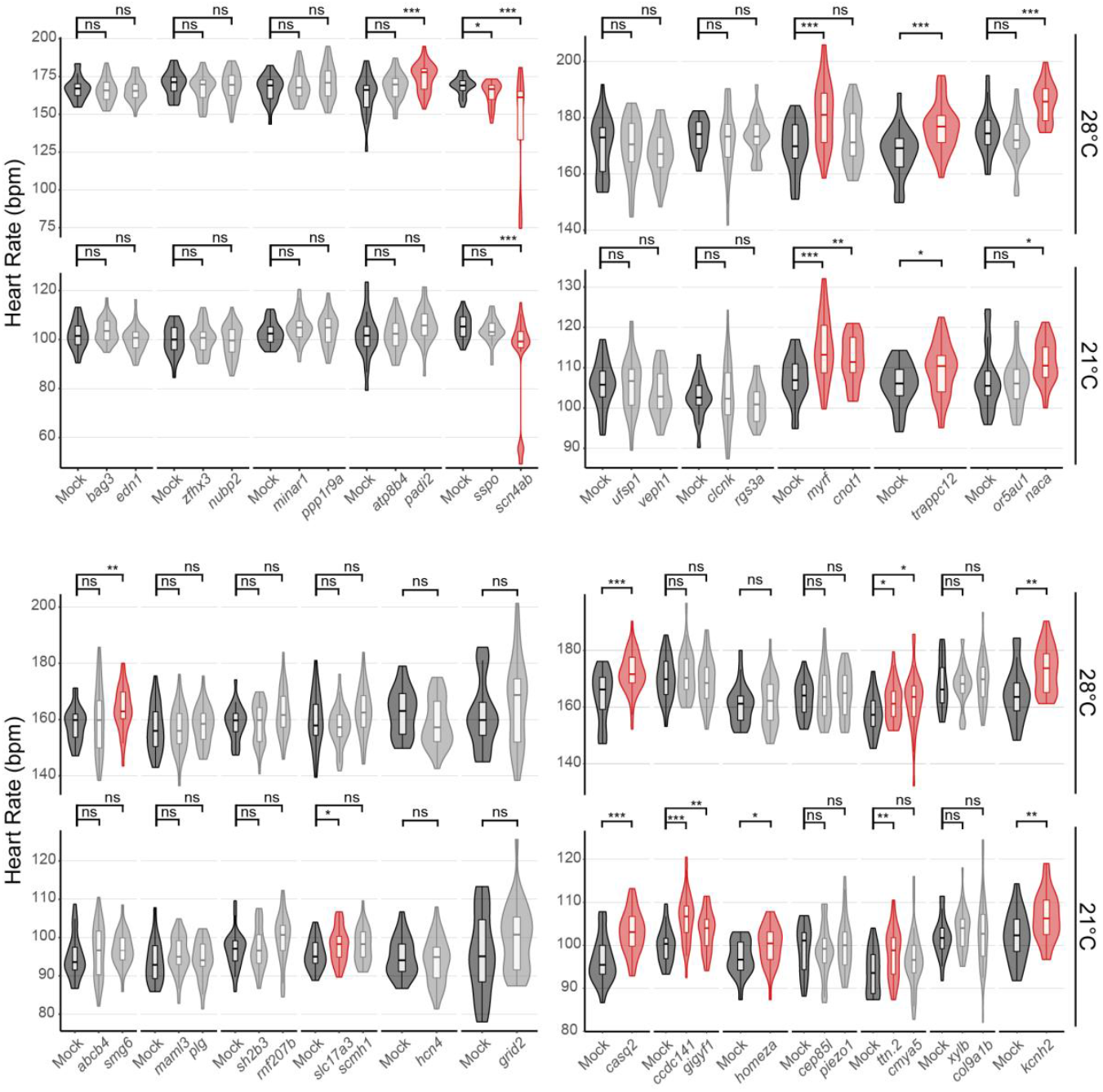
Comparative analysis of mean heart rates in targeted hGWAS gene selection. Heart rate measurements (beats per minute, bpm) of *GFP*-injected (Mock; dark grey) and corresponding sibling crispant embryos (4 dpf) at 21 and 28°C after developmental focusing (also see heatmap representation of the data in Fig 3A). Different experimental plates are represented by breaks on the x-axis. Significant differences in mean heart rates were determined between each crispant embryo group and its corresponding sibling control group by two-tailed Student’s t-test; \**p* < 0.05, \*\**p* < 0.01, \*\*\**p* < 0.001, ns (not significant). Red groups correspond to crispants showing significant heart rate phenotypes, and light grey groups correspond to crispants showing no significant heart rate phenotype. For biological replicates see Source Data Fig S4.

**Fig S5.**
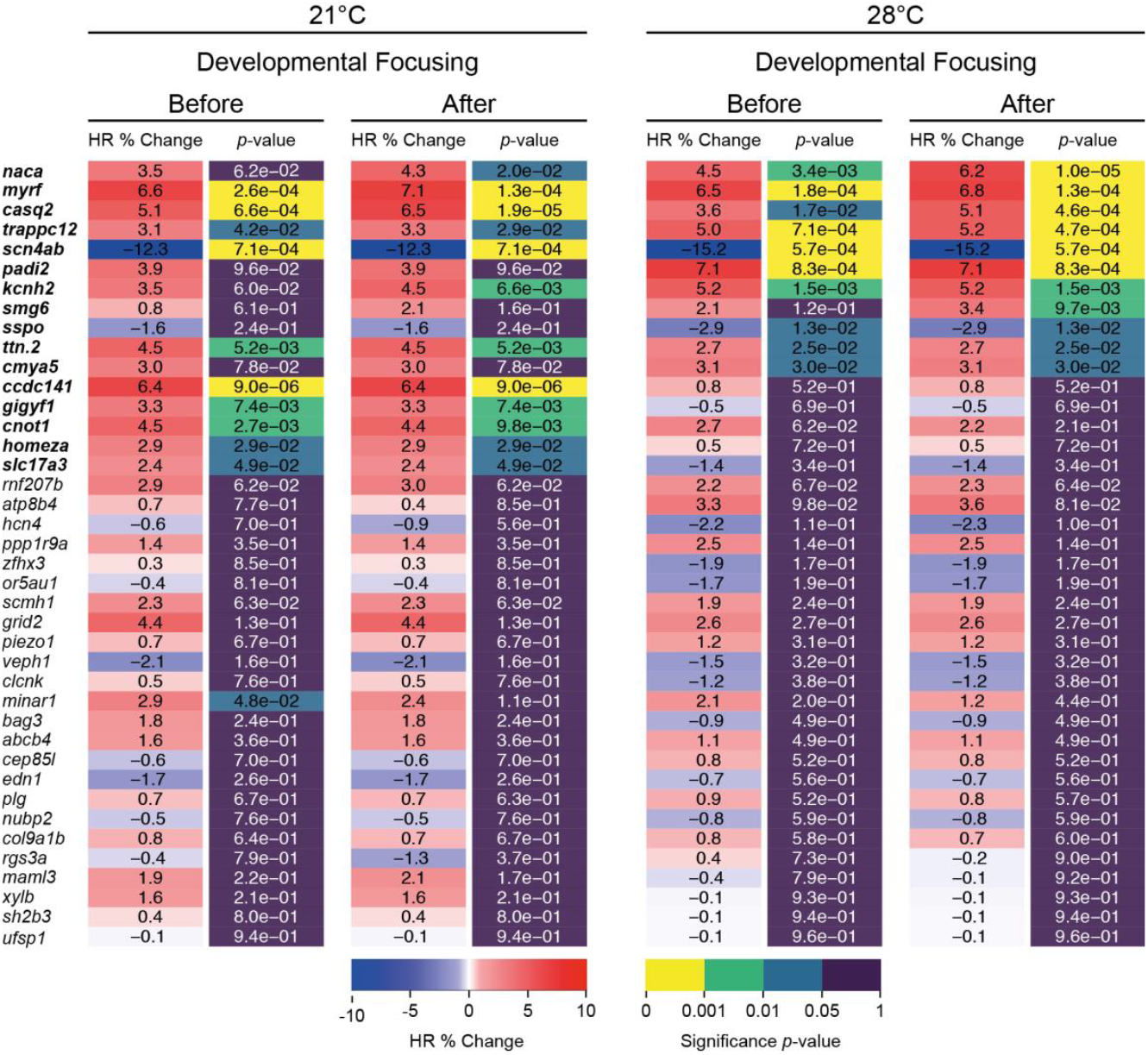
Developmental focusing does not alter analysis outcome of targeted hGWAS genes. Heatmap quantitative representation of the comparative heart rate analysis between each crispant embryo group and its corresponding control sibling group before and after developmental focusing; for each measured temperature, the percent change in mean heart rate (HR % Change) between crispants and their corresponding control siblings, flanked by the statistical significance (*p*-value) of the observed change, calculated by two-tailed Student’s t-test on the full distribution. Genes showing significantly different heart rate phenotypes are indicated in **bold**. For biological replicates see Source Data Fig S4.

**Movie S1. Moderate AV-block arrhythmia observed in medaka F0 *scn4ab* crispants and homozygous F2 mutants**

Side by side comparison of rhythmic heartbeat of *GFP*-injected (Mock; left) and arrhythmic *scn4ab* crispants (F0; middle) as well as homozygous mutants (F2; right) displaying 2:1 AV-block phenotype. Videos of medaka embryos (5 dpf) were acquired using a stereomicroscope under bright field illumination.

**Movie S2. Severe AV-block arrhythmia observed in medaka *scn4ab* crispants**

Side by side comparison of rhythmic heartbeat of *GFP*-injected (Mock; left) and arrhythmic *scn4ab* crispants displaying severe AV-block phenotype (right). Videos of medaka embryos (9 dpf) were acquired using a stereomicroscope under bright field illumination.

**Table S1.**
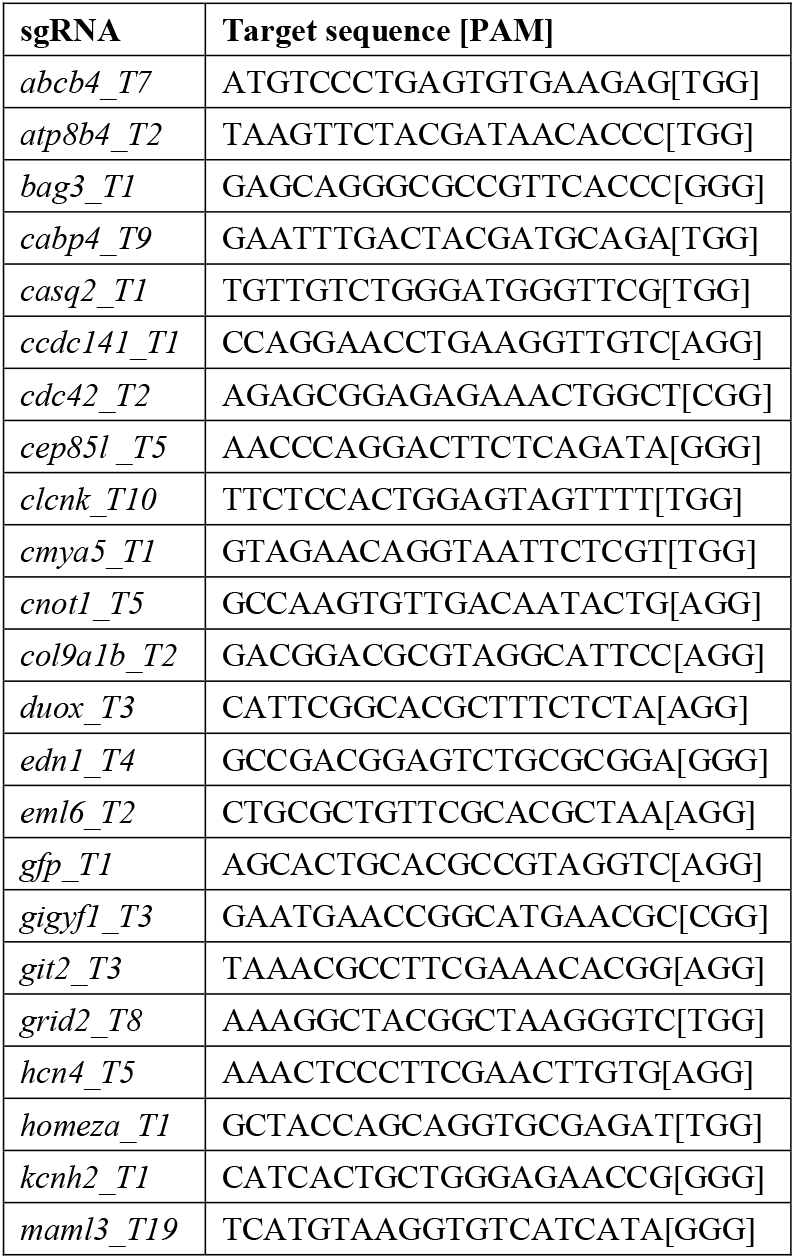
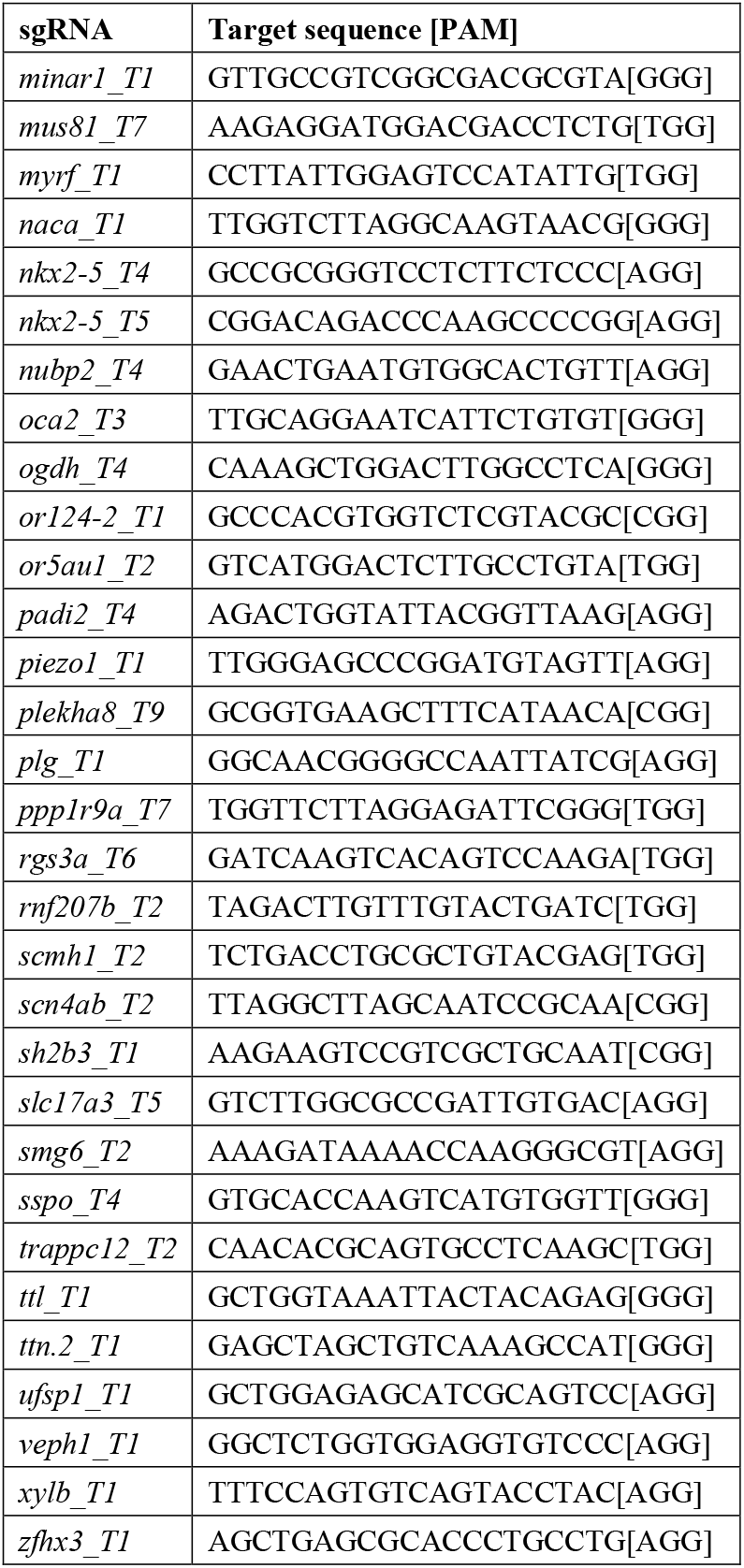
List of sgRNAs used. sgRNA target sites given in 5’-3’ direction, protospacer adjacent motif (PAM) in square brackets.

**Table S2.**
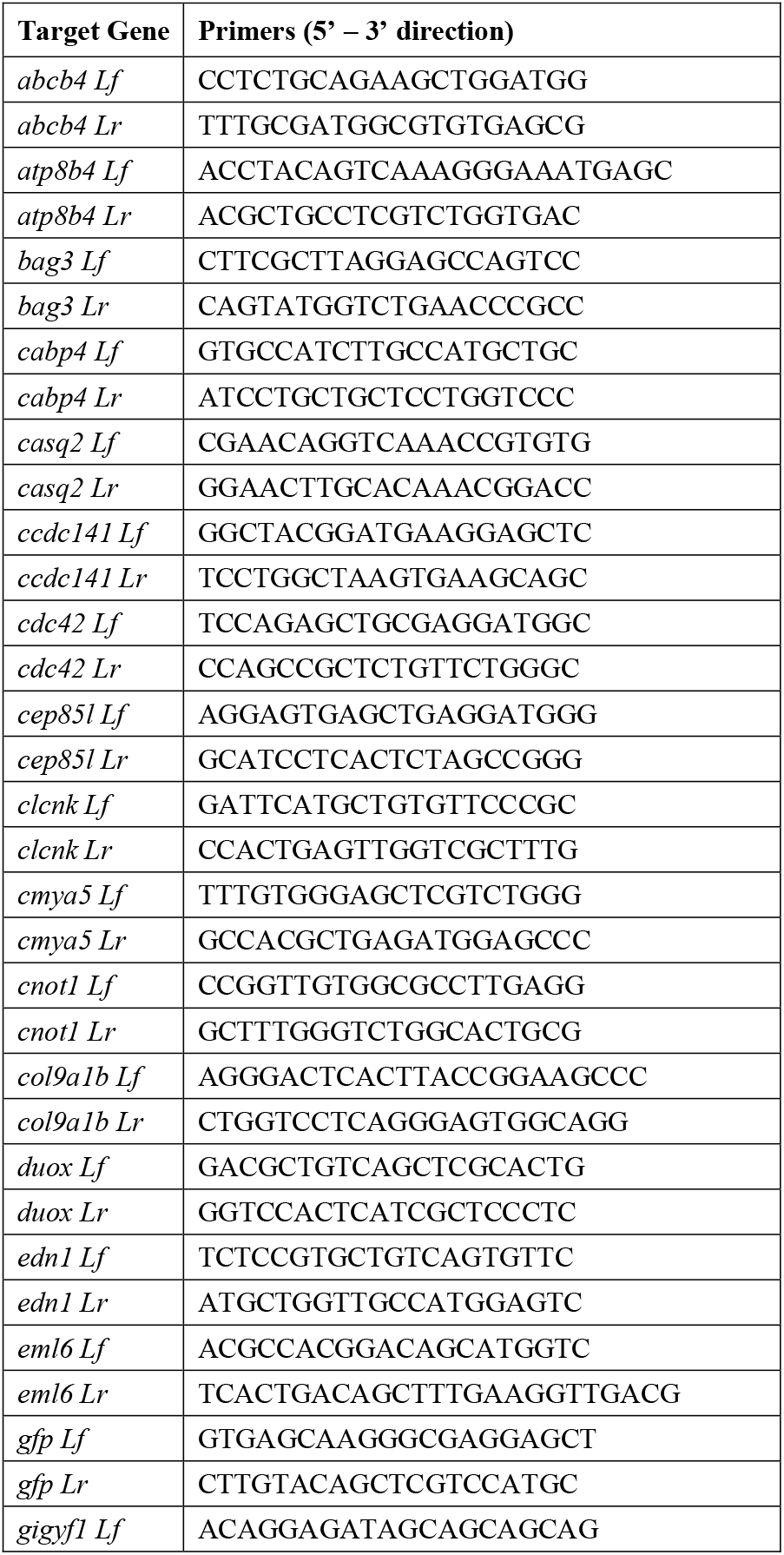
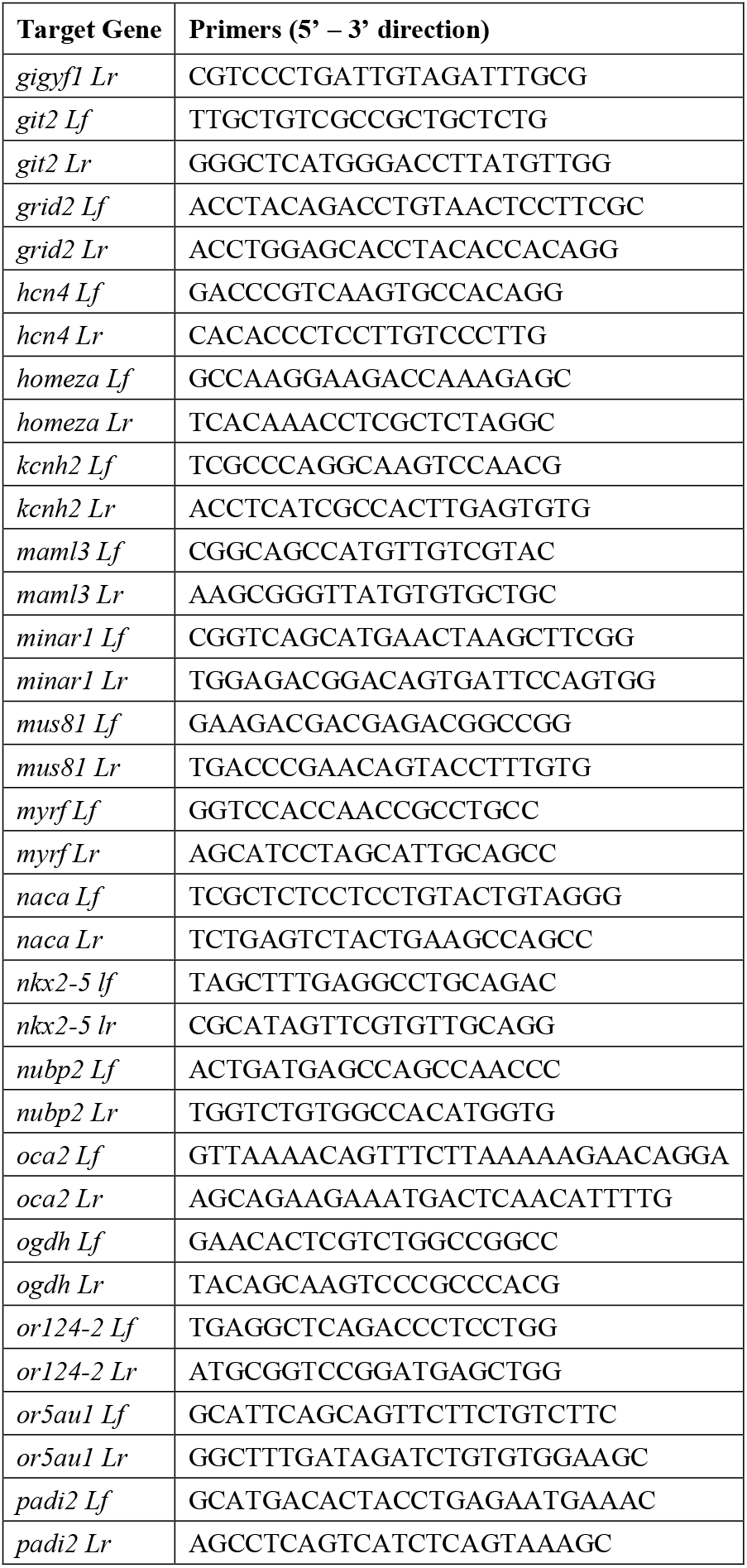
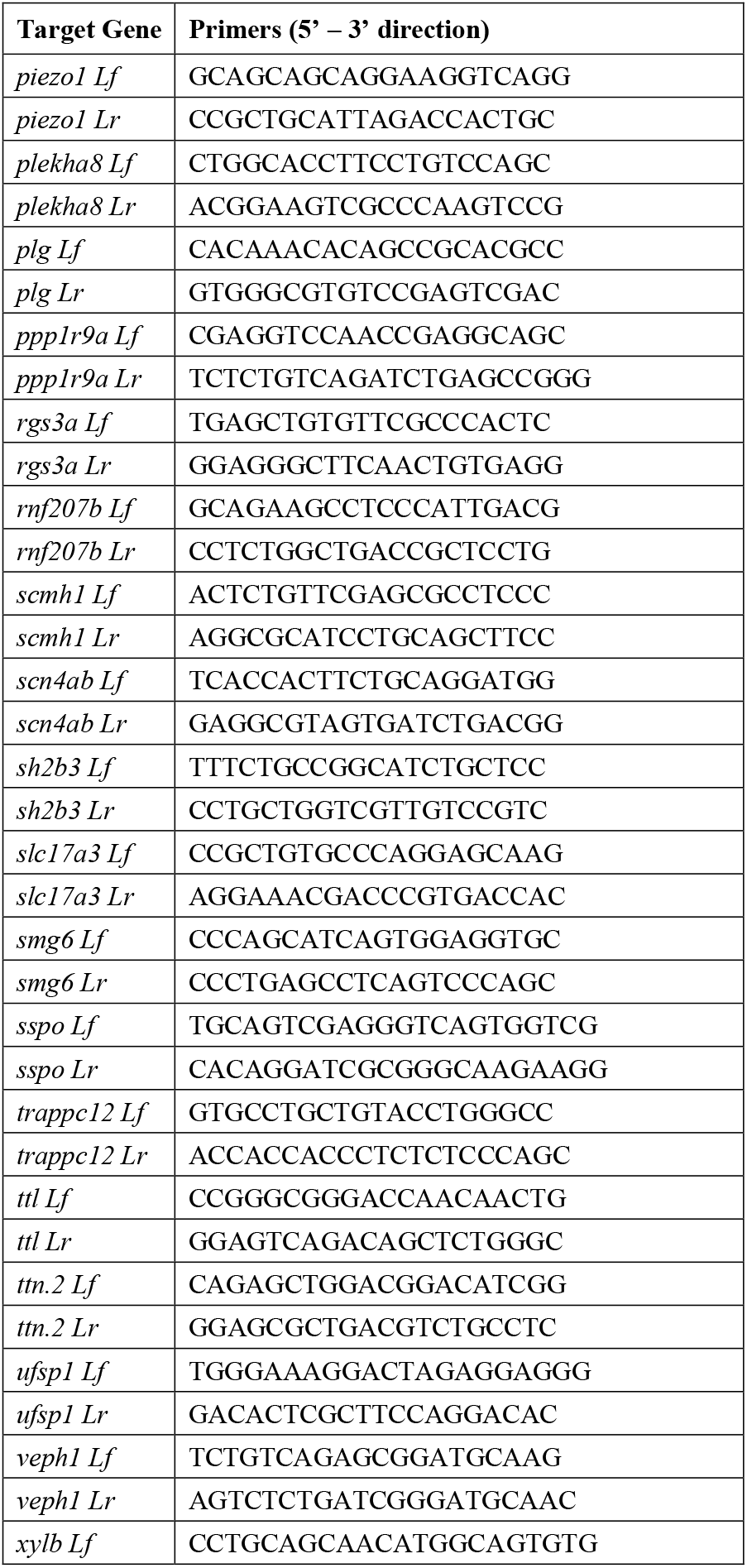
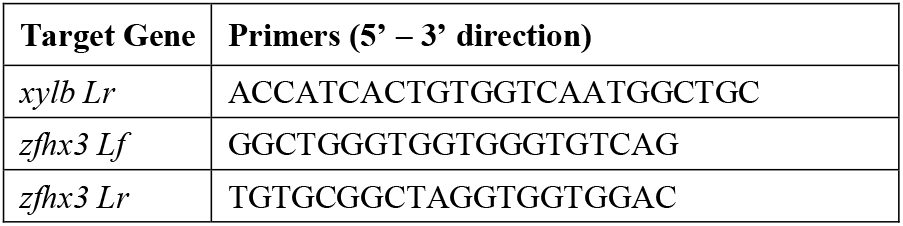
List of primers used for genotyping by PCR.

**Source Data Fig 1D. Biological replicates for Fig 1D**

Ns correspond to number of scored embryos per condition. before DF = before developmental focusing (i.e. all scored embryos). after DF = after developmental focusing (i.e. only scored embryos looking older than stage 28)

| Gene_sgRNA | Ns before DF |  | Ns after DF |  | Ns mock |  |
| --- | --- | --- | --- | --- | --- | --- |
|  | 21C | 28C | 21C | 28C | 21C | 28C |
| *nkx2-5_T4* | 34 | 34 | 31 | 31 | 22 | 23 |
| *oca2_T1* | 35 | 35 | 35 | 35 |  |  |

**Source Data Fig S2. Biological replicates for Fig S2 B-C**

Ns correspond to number of scored embryos per condition. before DF = before developmental focusing (i.e. all scored embryos). after DF = after developmental focusing (i.e. only scored embryos looking older than stage 28)

| Gene_sgRNA | Ns before DF |  | Ns after DF |  | Ns mock |  |
| --- | --- | --- | --- | --- | --- | --- |
|  | 21C | 28C | 21C | 28C | 21C | 28C |
| *nkx2-5_T4* | 34 | 36 | 31 | 33 | 19 | 23 |
| *nkx2-5_T5* | 33 | 33 | 31 | 31 | 22 | 21 |

**Source Data Fig S3. Biological replicates for Fig 2 and S3**

Ns correspond to number of scored embryos per condition. before DF = before developmental focusing (i.e. all scored embryos). after DF = after developmental focusing (i.e. only scored embryos looking older than stage 28)

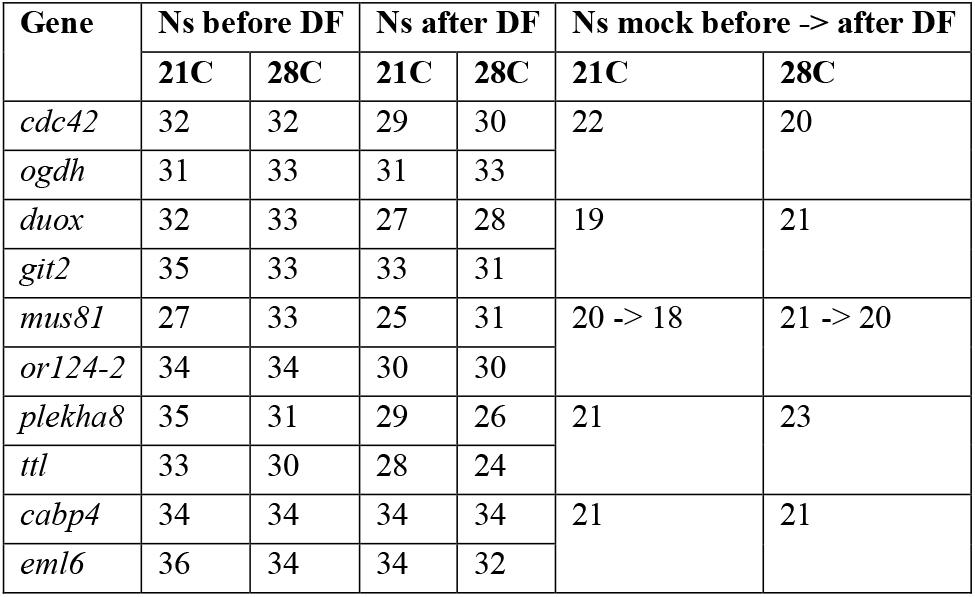

**Source Data Fig S4.** Biological replicates for Fig 3A, S4 and S5

Ns correspond to number of scored embryos per condition. before DF = before developmental focusing (i.e. all scored embryos). after DF = after developmental focusing (i.e. only scored embryos looking older than stage 28)

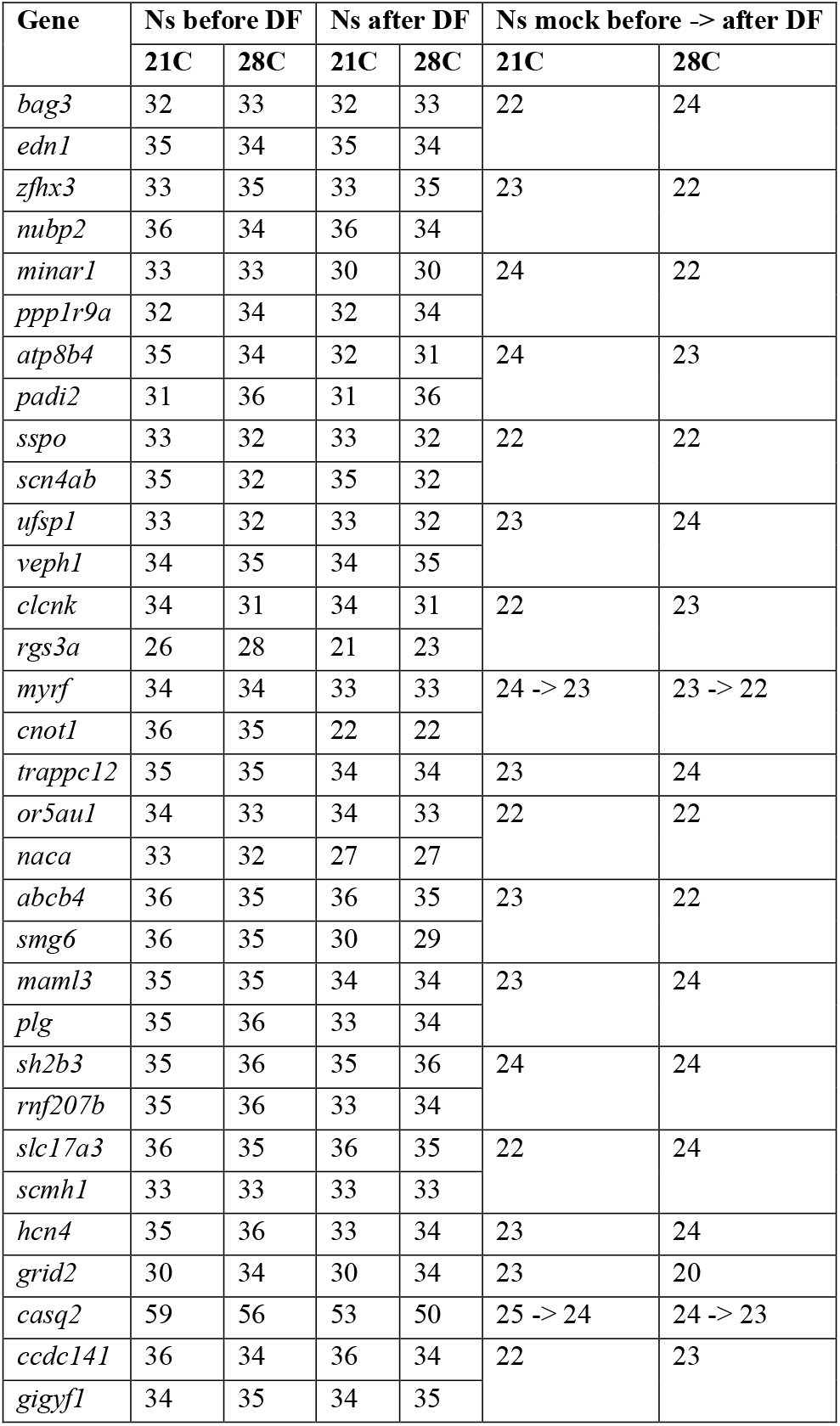
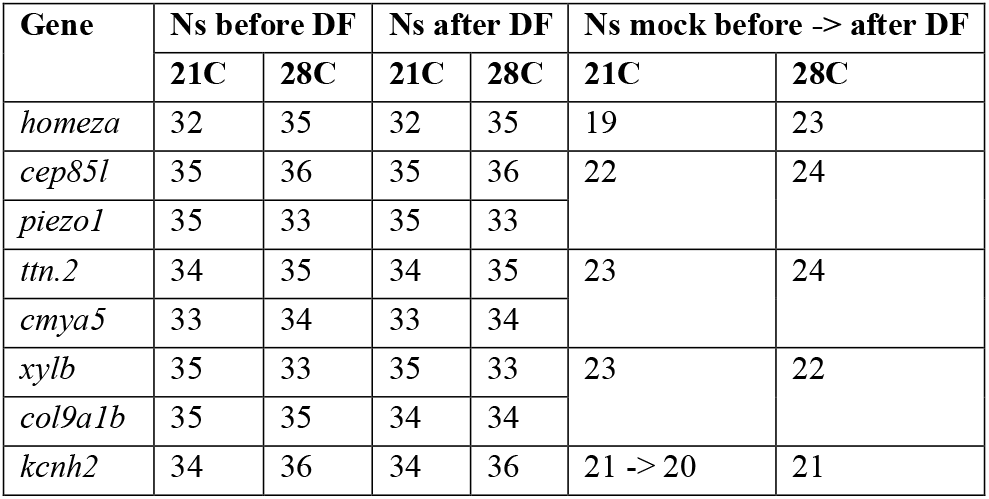

